# A suite of transgenic driver and reporter mouse lines with enhanced brain cell type targeting and functionality

**DOI:** 10.1101/224881

**Authors:** Tanya L. Daigle, Linda Madisen, Travis A. Hage, Matthew T. Valley, Ulf Knoblich, Rylan S. Larsen, Marc M. Takeno, Lawrence Huang, Hong Gu, Rachael Larsen, Maya Mills, Alice Bosma-Moody, La’Akea Siverts, Miranda Walker, Lucas T. Graybuck, Zizhen Yao, Olivia Fong, Emma Garren, Garreck Lenz, Mariya Chavarha, Julie Pendergraft, James Harrington, Karla E. Hirokawa, Julie A. Harris, Medea McGraw, Douglas R. Ollerenshaw, Kimberly Smith, Christopher A. Baker, Jonathan T. Ting, Susan M. Sunkin, Jerome Lecoq, Michael Z. Lin, Edward S. Boyden, Gabe J. Murphy, Nuno da Costa, Jack Waters, Lu Li, Bosiljka Tasic, Hongkui Zeng

**Author notes:** These authors contributed equally (Tanya L. Daigle, Linda Madisen).

## Abstract

Modern genetic approaches are powerful in providing access to diverse types of neurons within the mammalian brain and greatly facilitating the study of their function. We here report a large set of driver and reporter transgenic mouse lines, including 23 new driver lines targeting a variety of cortical and subcortical cell populations and 26 new reporter lines expressing an array of molecular tools. In particular, we describe the TIGRE2.0 transgenic platform and introduce Cre-dependent reporter lines that enable optical physiology, optogenetics, and sparse labeling of genetically-defined cell populations. TIGRE2.0 reporters broke the barrier in transgene expression level of single-copy targeted-insertion transgenesis in a wide range of neuronal types, along with additional advantage of a simplified breeding strategy compared to our first-generation TIGRE lines. These novel transgenic lines greatly expand the repertoire of high-precision genetic tools available to effectively identify, monitor, and manipulate distinct cell types in the mouse brain.

## INTRODUCTION

The nervous system is composed of a myriad of cell types in a highly organized and coordinated manner to produce function (Ascoli et al., 2008; Markram et al., 2004; Masland, 2004; Nelson et al., 2006; Somogyi and Klausberger, 2005; Zeng and Sanes, 2017). Neuronal cell types have extremely diverse molecular, morphological and physiological properties, and are interconnected in an intricate and precise fashion to form multiple levels of neural circuits that enable nervous system-specific functions, such as sensation, awareness, behavior, emotion, and cognition. Large-scale efforts have been underway to systematically profile these properties at the single cell level and develop cell type classification schemes (Zeng and Sanes, 2017), to build connectomes at micro-, meso- and macroscopic levels (Lichtman and Denk, 2011), and to probe neuronal activities in behaving animals (Peron et al., 2015; Yang and Yuste, 2017). All these levels of investigation are necessary to obtain fundamental understanding of how the nervous system works. And central to all these investigations is our ability to gain access to individual cell types, to observe and manipulate them, to perform circuit dissection and to connect information gathered at gene, cell and network levels (Huang and Zeng, 2013; Luo et al., 2008).

Cell type-specific genetic tools have grown into a powerful and dominant approach for studies of brain function via targeted expression of a variety of molecular sensors and probes in different cell types. In the mouse, the most advanced mammalian genetic model organism, a binary system based on Cre recombinase and its recognition sites (Lox sites) has been applied in nearly all areas of brain research, with a large number of Cre driver transgenic mouse lines now available to target specific populations of cells in various brain regions (Gerfen et al., 2013;

Gong et al., 2007; Harris et al., 2014; Madisen et al., 2010; Taniguchi et al., 2011). This combined with other recombinases and transcriptional regulation systems (Dymecki et al., 2010; Huang and Zeng, 2013; Madisen et al., 2015) in transgenic mouse lines and/or recombinant viral vectors has allowed increasingly more refined and flexible control of specific cell types. Recent rapid advances in single-cell transcriptional profiling (Ecker et al., 2017; Poulin et al., 2016;

Zeng and Sanes, 2017) have yielded novel cell type-specific marker genes and this, coupled with the continued development of new and better genetically encoded molecular sensors and probes, offers an ever wider array of options to fuel the genetic tool generation pipeline. Overall, the creation of new cell type-specific genetic tools holds tremendous promise in enabling better labeling, monitoring and manipulation of cell types as well as neural circuits and, ultimately, detailed mechanistic insight on brain function. At the same time, the demand for genetic tools with better specificity and performance still far exceeds their current availability, and much work is needed to convert promise into reality.

Cell type-specific genetic toolkits are usually designed as binary systems: they consist of two components – a driver and a reporter (can also be called a responder or effector) (Huang and Zeng, 2013). Usually, drivers and reporters are created separately, so that they can be combined to enable multiplicative growth of applications from a linear increase of the total number of lines. Either of the two components can be delivered in the form of a stably integrated transgene or a recombinant viral vector. Furthermore, enhanced specificity can be obtained by intersectional approaches that combine two or more binary systems. In the past decade, in addition to the creation of Cre, Flp, and other driver lines, we and others have built a range of Cre reporter lines and intersectional reporter lines based on the ubiquitously permissive Rosa26 locus (Dymecki et al., 2010; He et al., 2016; Madisen et al., 2012; Madisen et al., 2010; Muzumdar et al., 2007; Soriano, 1999). We also explored another permissive genomic locus, TIGRE, to establish a series of intersectional reporter lines controlled by Cre and a tetracycline-regulated transcriptional transactivator, tTA (Madisen et al., 2015; Zeng et al., 2008). We found that these TIGRE-based lines afforded substantially higher reporter and effector gene expression compared to the Rosa26-based lines, likely due to tTA/TRE-mediated transcriptional amplification (Madisen et al., 2015). This is a critical improvement as sufficiently high-level expression of many molecular tools, such as fluorescent proteins, calcium indicators and channelrhodopsins, is needed to allow full-extent visualization of fine neuronal structures, and effective reporting or manipulation of neuronal activities *in vivo* in a minimally invasive fashion.

In this paper, we report a large and diverse set of new cell type-specific transgenic mouse lines, including 23 new driver lines (15 Cre drivers and 8 Flp drivers) and 26 new reporter lines. The reporter lines express a variety of molecular tools that will enable a wide range of applications, such as synaptic or nuclear targeted fluorescent proteins, an electron microscopy tag, genetically encoded calcium indicators (GECIs), genetically encoded voltage indicators (GEVIs) and new channelrhodopsin variants. In particular, we have developed a new transgene design in the TIGRE locus, in which a Cre-dependent tTA-expressing transgene cassette was co-integrated into TIGRE along with the Cre and tTA-dependent reporter gene cassette. This approach effectively converts the TIGRE reporter lines into simple Cre-reporters while retaining the tTA-mediated transcriptional amplification. We name these second-generation TIGRE lines TIGRE2.0, and our previous Cre/tTA-dependent TIGRE lines TIGRE1.0. TIGRE2.0 lines simplify breeding strategies and expand the repertoire of cell types with strong reporter gene expression to nearly all the cell types we have examined. Quantitative gene expression analysis shows that TIGRE2.0 reporters have reached a transgene expression level comparable to (or higher than) that of a strong promoter-driven recombinant adeno-associated virus (AAV) vectors, breaking a significant barrier in single-copy targeted-insertion transgenesis. Importantly, we demonstrate superior functionality of many new transgenic lines, and also disclose side effects observed in animals with extremely high-level and widespread transgene expression in some circumstances. Experiences gained and lessons learned in the extensive study reported here will be invaluable for the utilization and further development of optimal genetic targeting strategies.

## RESULTS

The transgenic mouse program at the Allen Institute for Brain Science has generated transgenic mouse lines for cell type-specific labeling and manipulation for the past 10 years (Harris et al., 2014; Madisen et al., 2015; Madisen et al., 2012; Madisen et al., 2010; Tasic et al., 2016; Wang et al., 2017; Zariwala et al., 2012). A total of 56 driver lines and 54 reporter lines have been generated. Transgene expression patterns throughout the entire mouse brain for vast majority of these lines have been systematically characterized using our RNA *in situ* hybridization (ISH) pipeline in a standardized manner, and all the data are available at the brain-map.org portal (http://connectivity.brain-map.org/transgenic; http://www.alleninstitute.org/what-we-do/brain-science/research/products-tools/). To provide a comprehensive and centralized overview of the program, all transgenic lines generated in the Allen Institute are summarized in Table 1 for driver lines and Table 2 for reporter lines. Most of these lines have been deposited to the Jackson Laboratory, and they have been widely utilized in the broad research community.

**Table 1.**
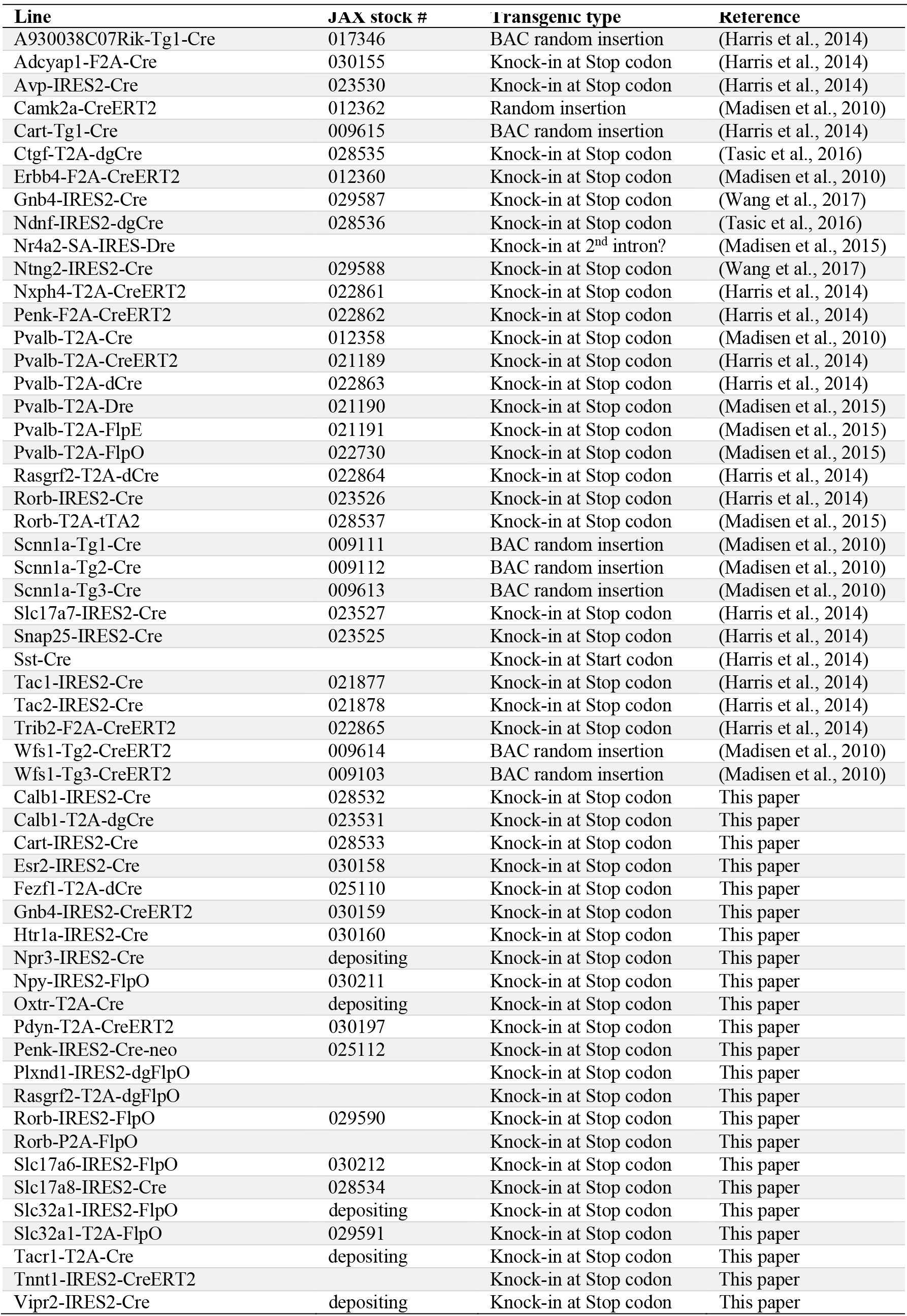
Driver lines generated by the Allen Institute for Brain Science

**Table 2.**
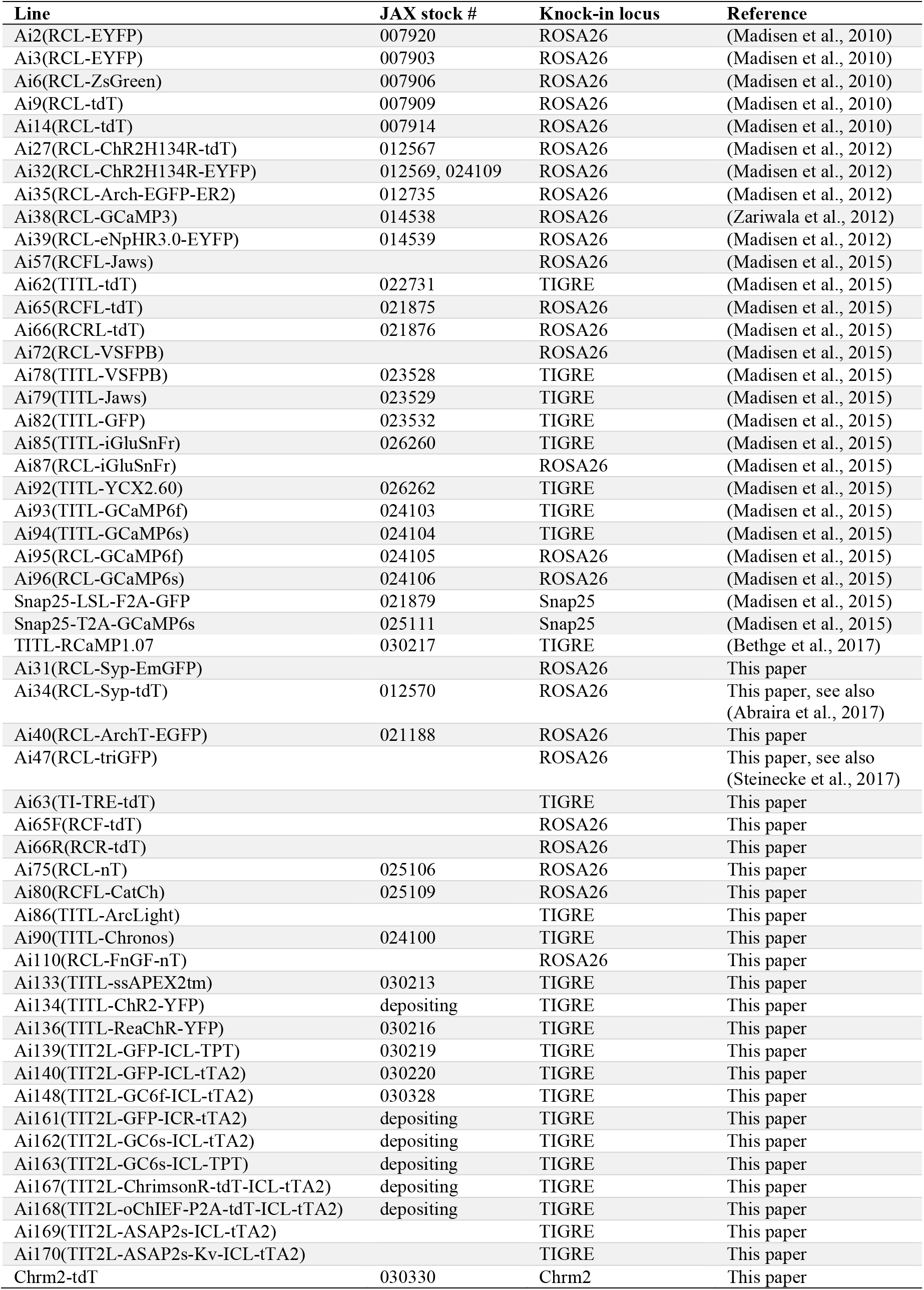
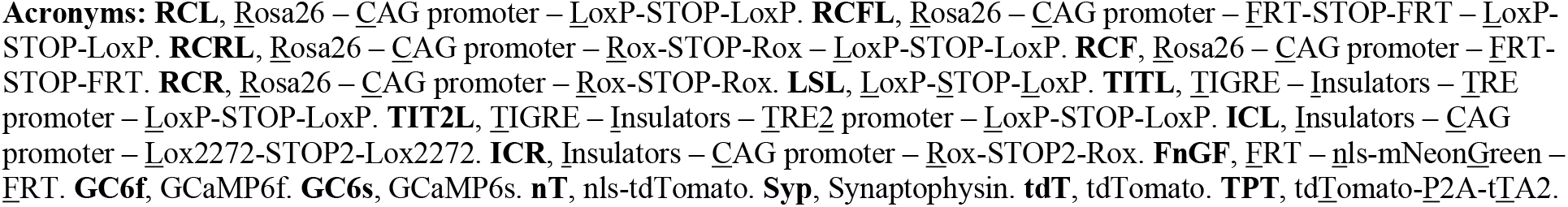
Reporter lines generated by the Allen Institute for Brain Science

Out of these 110 lines, 33 driver lines and 28 reporter lines have been described in previous publications, whereas the remaining 23 driver lines and 26 reporter lines are formally reported for the first time in this paper.

### New Cre and Flp driver lines targeting a variety of cell populations in cortical and subcortical areas

The 23 new driver lines (Table 1) were created by inserting a Cre or FlpO driver gene into the C-terminus of an endogenous gene via homologous recombination, using either conventional or CRISPR/Cas9-mediated approaches, with driver gene expression mediated by either an internal ribosome entry site (IRES, variant IRES2 in particular) or a 2A peptide (T2A or P2A) sequence. Out of these, 15 lines express variants of Cre, including Cre itself, tamoxifen-inducible CreERT2, trimethoprim (TMP)-inducible dCre (stands for DHFR-Cre) or dgCre (stands for DHFR-GFP-Cre) (Sando et al., 2013), and 8 lines express FlpO or its TMP-inducible version dgFlpO (DHFR-GFP-FlpO). The GFP part present in dgCre and dgFlpO lines exhibits no visible fluorescence with or without TMP induction.

Target genes chosen for making Cre and FlpO driver lines to supplement existing lines include cortical layer-specific genes (*e.g., Calb1, Htr1a, Npr3, Plxnd1, Rasgrf2*, and *Rorb*), neuropeptide and receptor genes (e.g., *Cart, Esr2, Npy, Oxtr, Pdyn, Penk, Tacr1*, and *Vipr2*), genes targeting specific cell populations (e.g., *Fezf1* for ventromedial hypothalamus, *Gnb4* for claustrum, and *Tnnt1* for thalamus), and pan-GABAergic (*Slc32a1*, also known as *Vgat*) or glutamatergic (*Slc17a6*, also known as *Vglut2*, and *Slc17a8*, also known as *Vglut3*) genes. Expression patterns of the driver genes as well as the tdTomato reporter gene in corresponding reporter lines, Ai14 for Cre and Ai65F (see below) for FlpO were examined by ISH (Fig. 1 and http://connectivity.brain-map.org/transgenic).

**Figure 1.**
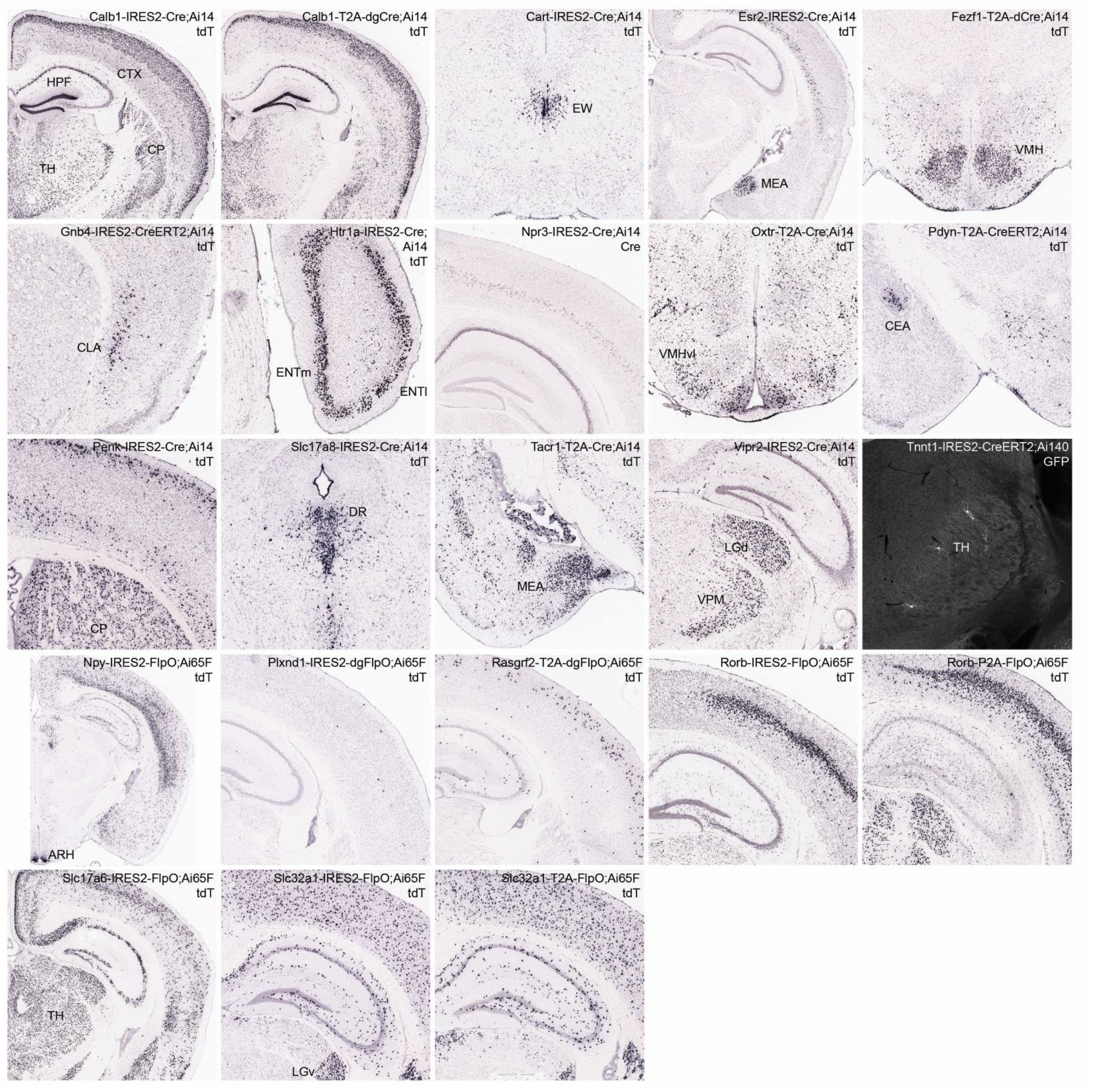
New Cre and Flp driver lines targeting specific cell populations in the brain. Selected images from RNA *in situ* hybridization (ISH) characterization of all new driver lines using either tdTomato reporter probe or Cre probe as indicated on each panel. Full ISH datasets are viewable at http://connectivity.brain-map.org/transgenic. The Cre-dependent reporter Ai14 was used to examine all the Cre driver lines, and the Flp-dependent reporter Ai65F (see Fig. S1) was used to examine all the FlpO driver lines. The only exception is Tnnt1-IRES2-CreERT2, which was crossed to a new TIGRE2.0 EGFP-expressing Cre reporter line Ai140 and TissueCyte imaging shows extremely sparse GFP labeling of thalamic neurons (after 1-day tamoxifen induction at P10). ARH, arcuate hypothalamic nucleus. CEA, central amygdalar nucleus. CLA, claustrum. CP, caudoputamen. CTX, cortex. DR, dorsal nucleus raphe. ENTl, entorhinal area, lateral part. ENTm, entorhinal area, medial part. EW, Edinger-Westphal nucleus. HPF, hippocampal formation. LGd, dorsal part of the lateral geniculate complex. LGv, ventral part of the lateral geniculate complex. MEA, medial amygdalar nucleus. TH, thalamus. VMH, ventromedial hypothalamic nucleus. VPM, ventral posteromedial nucleus of the thalamus.

We find that the expression patterns of all driver transgenes largely recapitulate the adult (P56) expression patterns of their endogenous target genes. On the other hand, the expression patterns of Cre- and Flp-reporters are often more widespread, likely due to the cumulative recombination from the transient expression of the driver genes in developmental or adult stages. An observed trend is that IRES-mediated drivers tend to drive more faithful expression of the reporters whereas 2A-mediated drivers often lead to more widespread expression of the transgenic reporters. Also, dCre and dgCre (from Calb1-T2A-dgCre and Fezf1-T2A-dCre lines) can mediate efficient recombination as do Cre (from Calb1-IRES2-Cre, Cart-IRES2-Cre, Esr2-IRES2-Cre, Htr1a-IRES2-Cre, Npr3-IRES2-Cre, Oxtr-T2A-Cre, Penk-IRES2-Cre-neo, Slc17a8-IRES2-Cre, Tacr1-T2A-Cre and Vipr2-IRES2-Cre) and FlpO (from Npy-IRES2-FlpO, Rorb-IRES2-FlpO, Rorb-P2A-FlpO, Slc17a6-IRES2-FlpO, Slc32a1-IRES2-FlpO and Slc32a1-T2A-FlpO), whereas CreERT2 (from Gnb4-IRES2-CreERT2, Pdyn-T2A-CreERT2 and Tnnt1-IRES2-CreERT2) and dgFlpO (from Plxnd1-IRES2-dgFlpO and Rasgrf2-T2A-dgFlpO) tend to have less efficient recombination as indicated by the sparser activation of the reporters.

### New Rosa26 and TIGRE1.0 reporter lines with driver-dependent expression of new genetic tools

We first summarize a set of Rosa26-based Cre reporter lines we generated but have not formally reported previously (Table 2 and Fig. S1). These include Ai31 (expressing synapse-localized Synaptophysin-EmGFP), Ai34 (expressing synapse-localized Synaptophysin-tdTomato) (Abraira et al., 2017), Ai40 (expressing archeorhodopsin ArchT-GFP) (Han et al., 2011), Ai47 (expressing 3 tandemly-linked GFP molecules EmGFP-T2A-TagGFP2-P2A-hrGFP) (Steinecke et al., 2017), Ai75 (expressing nuclear-localized nls-tdTomato), Ai80 (expressing channelrhodopsin CatCh) (Kleinlogel et al., 2011), Ai86 (expressing voltage sensor ArcLight) (Jin et al., 2012), and Ai110 (a Cre/Flp dual reporter expressing nuclear-localized nls-mNeonGreen when only Cre is present and nls-tdTomato when both Cre and Flp are present). All of these lines display appropriate reporter gene expression when crossed to various Cre lines (Fig. S2A-D).

We also generated a knock-in Chrm2-tdT line, in which tdTomato was fused to the C-terminus of the endogenous muscarinic acetylcholine receptor 2 gene (*Chrm2*) (Fig. S1). Immunohistochemical staining of the CHRM2 protein can be used to delineate cytoarchitectonic boundaries of various cortical areas (such as primary visual cortex and primary somatosensory cortex) (Ji et al., 2015). We find that such delineation is preserved by native fluorescence of the CHRM2-tdTomato fusion protein in Chrm2-tdT mice (Fig. S2E); thus this line can be used to visualize cortical area boundaries *in vivo*.

We created a Flp-reporter line, Ai65F, and a Dre-reporter line, Ai66R, (Fig. S1), not de novo, but by performing EIIa-Cre mediated germline deletion of the Lox-STOP-Lox (LSL) cassette from our previously reported, Rosa26 locus-based Cre/Flp double reporter line Ai65, and Cre/Dre double reporter line Ai66, respectively (Madisen et al., 2015). Using the same approach, we created a tTA-reporter line Ai63 (Fig. S1) from the TIGRE locus-based Cre/tTA double reporter line Ai62 (Madisen et al., 2015). Expected specificity for their respective driver lines was observed (Fig. 1 and Fig. S2F).

Following the successful targeting of the TIGRE locus in generating a battery of Cre/tTA double reporter lines expressing a variety of tools at very high levels (Madisen et al., 2015), we generated several additional lines expressing new tools in a similar manner. These include Ai90 (expressing channelrhodopsin Chronos-GFP) (Klapoetke et al., 2014), Ai133 (expressing an electron microscopy tag ssAPEX2tm) (Lam et al., 2015), Ai134 (expressing channelrhodopsin ChR2(H134R)-YFP), and Ai136 (expressing channelrhodopsin ReaChR-YFP) (Lin et al., 2013) (Fig. 2A). As shown before (Madisen et al., 2015), for the three new channelrhodopsin expressing lines (Ai90, Ai134 and Ai136), a triple transgenic cross (Cre x tTA x reporter) is needed to enable reporter gene expression. We found that the use of a moderately expressing tTA driver line, ROSA26-ZtTA (Li et al., 2010), led to good expression of all three channelrhodopsin proteins when combined with cortical layer 4- or 6-specific Cre lines (Scnn1a-Tg3-Cre or Ntsr1-Cre_GN220) (Fig. 2B). On the other hand, the use of a strong tTA driver, Camk2a-tTA, led to fluorescent aggregates in cells and/or aberrant morphology of labeled cells (data not shown), indicating adverse effects of overexpression of these opsin molecules. Thus, we advise against the use of Camk2a-tTA in these cases.

**Figure 2.**
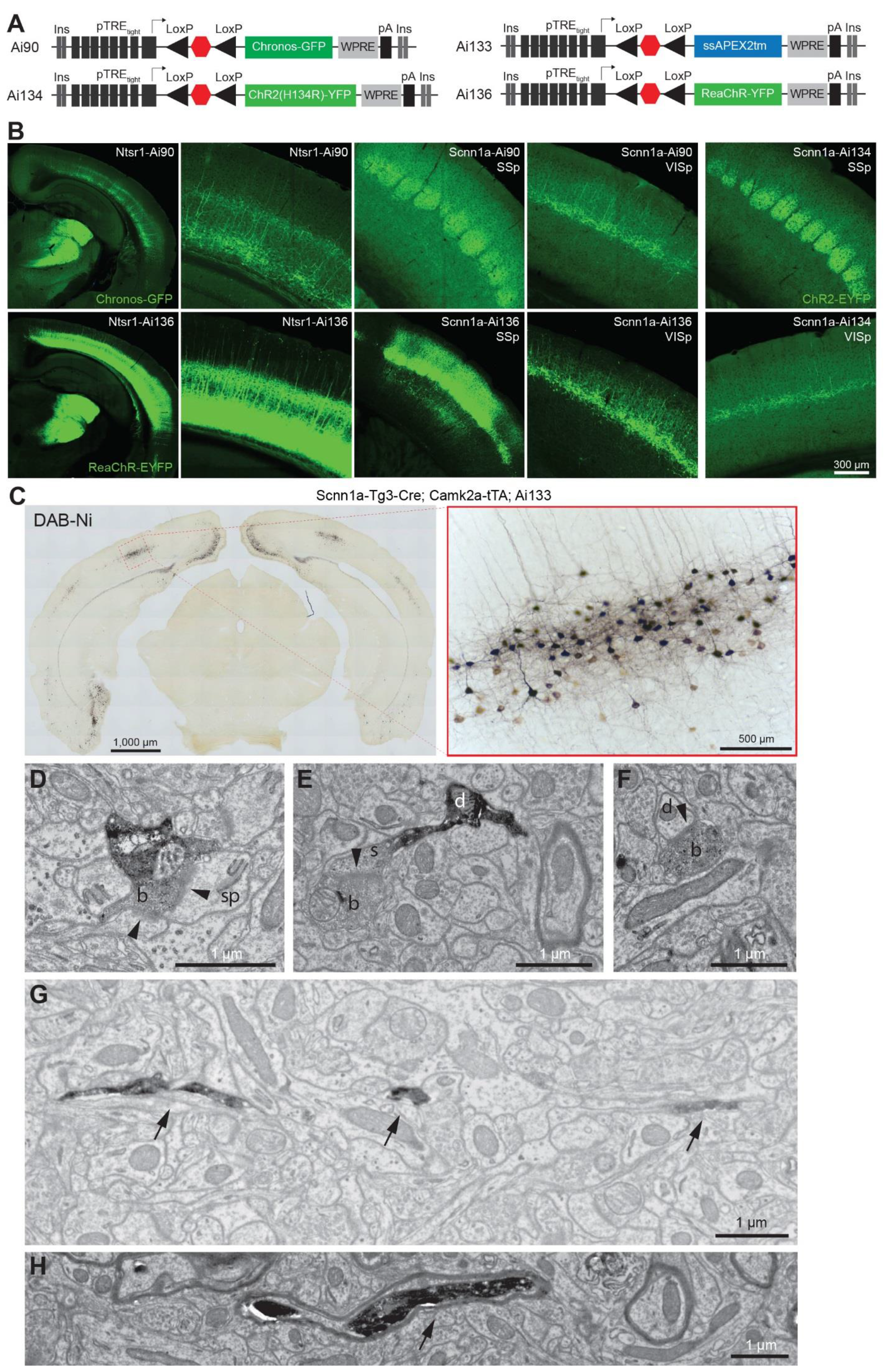
New TIGRE1.0 reporter lines for light-induced activation and ultrastructural labeling of genetically-defined cell types. (**A**) Schematic diagram of four new Cre- and tTA-dependent TIGRE1.0 reporter lines. Channelrhodopsin variants Chronos (Ai90), ChR2(H134R) (Ai134) and ReaChR (Ai136) were directly fused to GFP or YFP. Ascorbate peroxidase APEX2 was tagged with a T-cell CD2 antigen signal sequence (ss) and a transmembrane targeting sequence (tm). (**B**) Robust Cre-dependent expression of channelrhodopsin variants in cortical layer 4 and 6 neurons. Layer 6 expression is shown for Chronos in Ntsr1-Cre_GN220;ROSA26-ZtTA;Ai90 mice and for ReaChR in Ntsr1-Cre_GN220;ROSA26-ZtTA;Ai136 mice. Layer 4 expression is shown for Chronos in Scnn1a-Tg3-Cre;ROSA26-ZtTA;Ai90 mice, for ChR2 in Scnn1a-Tg3-Cre;ROSA26-ZtTA;Ai134 mice, and for ReaChR in Scnn1a-Tg3-Cre;ROSA26-ZtTA;Ai136 mice. (**C**) APEX2 activity revealed by DAB-Ni staining on sections from an Scnn1a-Tg3-Cre;Camk2a-tTA;Ai133 animal. Bright-field images reveal staining within somatic, dendritic and axonal compartments of neurons located in several cortical regions, including layer 4 of the primary visual cortex (20X image; right). (**D-F**) Electron micrographs show APEX2-dependent labeling of both pre- and post-synaptic terminals. Examples include synapses (arrowheads) between a labelled bouton (b) and an unlabeled spine (sp) (D), between a labelled bouton and a labeled spine (E), and between a labelled bouton and an unlabeled dendritic shaft (d) (F). (**G-H**) Electron micrographs show APEX-dependent labeling of fine, unmyelinated (G) or myelinated (H) axons (arrows).

The Ai133 reporter line (Fig. 2A) expresses a membrane-tagged ascorbate peroxidase APEX2 (Lam et al., 2015), which we found to be compatible with transcardial perfusion and fixation with glutaraldehyde – a necessary step towards obtaining ultrastructure preservation for electron microscopy (EM). Incubation of coronal slices from Scnn1a-Tg3-Cre;Camk2a-tTA;Ai133 animals with 3,3’-diaminobenzidine enhanced with nickel (DAB-Ni) showed specific peroxidase activity in the cortical layer 4 as expected (Fig. 2C). Processing DAB-Ni stained tissue with a reduced osmium tetroxide (ROTO) protocol provided strong EM contrast and identification of APEX2-positive cell processes (Fig. 2D-H).

### TIGRE2.0 reporter lines with enhanced expression

The first generation TIGRE (TIGRE1.0) lines rely on two driver lines for cell type-specific and high-level transgene expression (Madisen et al., 2015) (Fig. 3A). This triple transgenic design requires laborious breeding and is limited by the availability of only a few well-functioning tTA driver lines (e.g. Camk2a-tTA, ROSA26-ZtTA, and Rorb-T2A-tTA2). By using these select tTA lines, we previously observed robust transgene expression in cortical excitatory neurons and a subset of cortical interneurons or subcortical neurons, but poor expression in many other neuronal types (Madisen et al., 2015). To overcome these problems and to simplify breeding, we investigated ways to incorporate tTA-mediated transcriptional amplification into the TIGRE locus itself along with the reporter transgene.

**Figure 3.**
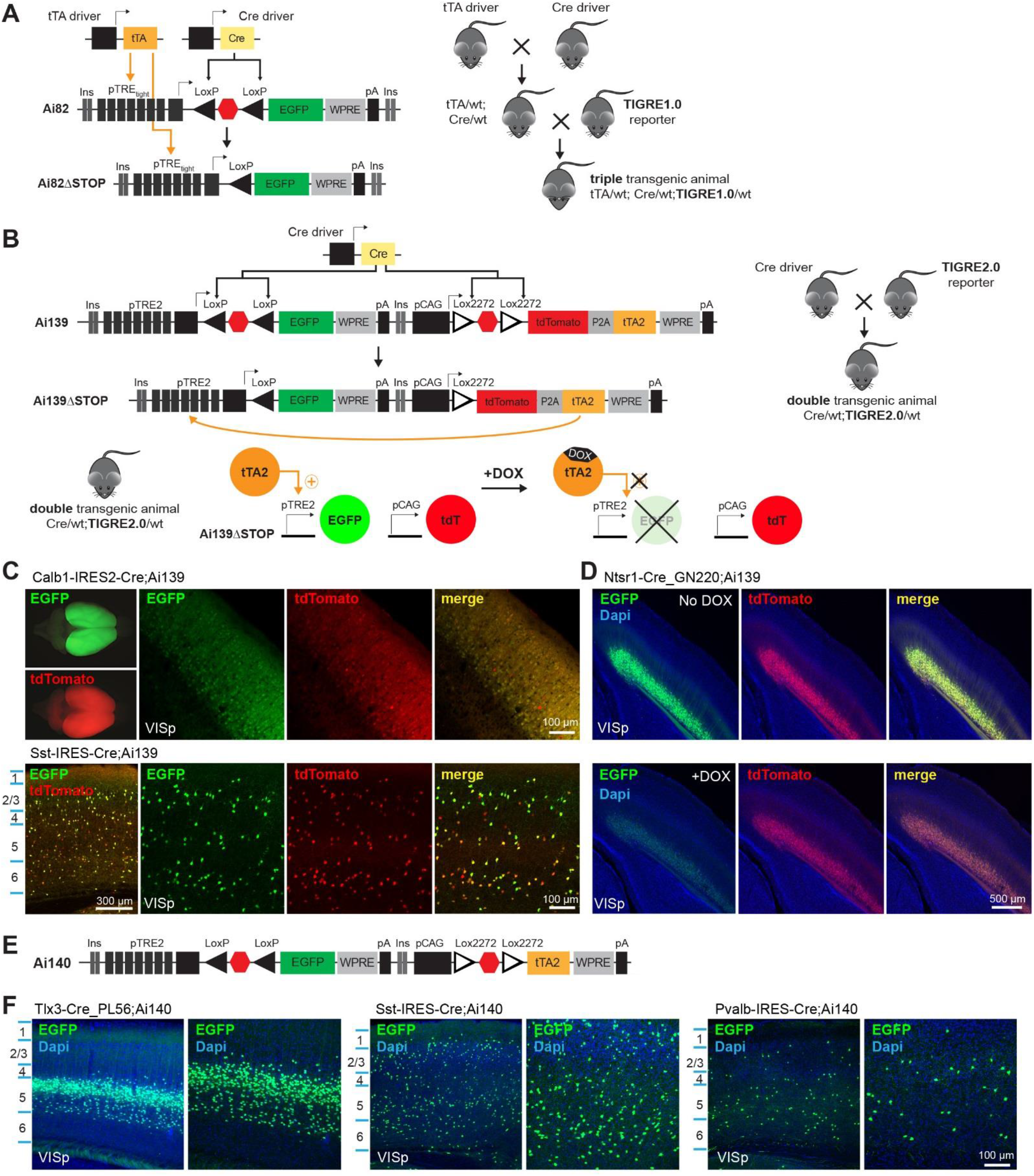
Incorporation of a tTA2-expressing unit within the TIGRE locus simplifies breeding and promotes strong transgene expression. See also Figure S3. (**A**) Schematic diagram of Cre- and tTA-dependent intersectional TIGRE1.0 reporter line design and generalized triple transgenic breeding scheme. (**B**) Schematic diagram of Cre-dependent TIGRE2.0 reporter line design and simplified double transgenic breeding scheme. In the first TIGRE2.0 reporter line, Ai139, Cre-mediated excision of the two LSL cassettes results in CAG promoter-driven co-expression of tdTomato and tTA2, the latter of which then promotes TRE2 promoter-driven EGFP expression. The tTA2 activity and subsequent TRE2 promoter-driven expression can be suppressed by giving doxycycline (DOX) to double transgenic animals. (**C**) Whole brain epifluorescence images (left panels) and representative confocal images of native EGFP and tdTomato fluorescence in the primary visual cortex (VISp) of Ai139 double transgenic animals. (**D**) Representative confocal images of native EGFP and tdTomato fluorescence in the VISp of Ntsr1-Cre;Ai139 mice (No Dox), or Ntsr1-Cre;Ai139 mice given DOX (+DOX). (**E-F**) Schematic diagram of Ai140, a Cre-dependent EGFP-expressing TIGRE2.0 reporter line (E), in which tTA2 is under the CAG promoter without the co-expression of tdTomato, and representative confocal images of native EGFP fluorescence in VISp from Ai140 double transgenic animals. Nuclei were stained with DAPI.

We first tried two approaches to co-express tTA and a GFP reporter gene, both under Cre control, via a 2A sequence or a bidirectional Bi-Tet promoter, with the hope that upon Cre-mediated recombination, a background level of tTA expression from the TRE or Bi-Tet promoter would initiate a self-amplifying cascade of expression of both tTA and the reporter gene (Reijmers et al., 2007). However, upon Cre transfection of targeted ES clones, we observed no GFP expression (data not shown). Since the TRE promoter we had used was the TRE-Tight version, we then replaced it with the original, ‘leakier’ tetO promoter (TRE2). In targeted ES clones for both 2A and Bi-Tet versions, we did observe GFP positive cells upon Cre transfection, however, the proportion of green cells was low (data not shown). We went ahead and generated a mouse line from the TRE2-LSL-GFP-2A-tTA2 clone (named Ai109). However, upon crossing Ai109 to 3 Cre lines, Pvalb-IRES-Cre, Nr5a1-Cre and Sst-IRES-Cre, we observed very few GFP-expressing cells in some but not all regions where Cre recombination would normally occur (data not shown). Thus we did not pursue this approach further.

The next approach we tried was to co-insert two complete transgene expressing units, separated by tandem HS4 insulators to mitigate potential interference between the two units, into the same TIGRE locus (Fig. 3B). The first unit contains TRE2 promoter-driven, LSL-controlled reporter gene (e.g., GFP), and the second unit contains CAG promoter-driven, LSL-controlled tTA2 gene (or tdTomato-P2A-tTA2). We named this approach TIGRE2.0. In the presence of Cre, both STOP signals would be excised, thereby allowing CAG promoter to drive tTA2 expression, which would in turn activate the reporter gene expression.

We developed two prototype TIGRE2.0 lines expressing GFP: Ai139(TIT2L-GFP-ICL-TPT) (stands for <u>T</u>IGRE-Insulators-<u>T</u>RE<u>2</u> promoter-<u>L</u>oxPStop1LoxP-<u>GFP</u>-Insulators-<u>C</u>AG promoter-<u>L</u>ox2272Stop2Lox2272-td<u>T</u>omato-<u>P</u>2A-t<u>T</u>A2), and Ai140(TIT2L-GFP-ICL-tTA2) (Fig. 3B,E). To determine if tTA2 expression from within the TIGRE locus would be sufficient, we crossed these two lines to several Cre lines and examined transgene expression levels in a variety of cell types. In all cases, we observed strong native GFP fluorescence throughout cortical and subcortical areas (Fig. 3C,F and Fig. S3) and remarkably, we also observed strong GFP fluorescence in *Pvalb*+ and *Sst*+ interneurons, which is a significant improvement over the TIGRE1.0 lines. Reporter expression could also be suppressed by administering doxycycline to double-transgenic animals (Fig. 3B,D).

### A suite of TIGRE2.0 reporter lines expressing functional probes in a wide range of cell types

With the above promising results, we proceeded to generate a set of TIGRE2.0 lines expressing other genetic tools for probing neuronal function. Those included genetically encoded calcium indicators (GECIs) GCaMP6f and GCaMP6s (Chen et al., 2013), new channelrhodopsin variants ChrimsonR (Klapoetke et al., 2014) and oChIEF (Ting et al., 2014), new genetically encoded voltage indicator (GEVI) ASAP2s (Chamberland et al., 2017; Yang et al., 2016), and a new Cre/Dre intersectional reporter line.

We generated three TIGRE2.0 reporter lines expressing GCaMP6f or GCaMP6s: Ai148(TIT2L-GC6f-ICL-tTA2), Ai162(TIT2L-GC6s-ICL-tTA2) and Ai163(TIT2L-GC6s-ICL-TPT) (Fig. 4A). To directly compare the cell type coverage of these new reporters relative to the previous Rosa26-based reporters, we generated triple transgenic mice containing both Ai14 and Ai148 along with one of six Cre lines (Fig. 4B). With a set of cortical layer-specific Cre lines (Fig. 4B, upper panels), we observed nearly complete overlap in the expression of GCaMP6f-positive and tdTomato-positive cells. With the 3 most common cortical interneuron-specific Cre lines (Fig. 4B; lower panels), GCaMP6f and tdTomato were found to be largely coexpressed at the single cell level, but we also detected cells expressing only tdTomato, indicating the difficulty of achieving tTA-dependent expression in interneuron populations is not fully overcome. This is particularly the case for *Pvalb+* interneurons, which had the lowest proportion of GCaMP6f-positive cells relative to the tdTomato-positive cells. Strong GCaMP6s expression was also observed when crossing Ai162 or Ai163 mice to a pan-cortical glutamatergic Cre line, Slc17a7-IRES2-Cre (Vglut1), or an interneuron Cre line, Pvalb-IRES-Cre (Fig. 4C). Quantification of the extent of GCaMP6f or GCaMP6s co-expression with tdTomato at the single cell level confirmed the above observations, and also revealed that GCaMP6s was expressed in a larger (in Ai162) or substantially larger (in Ai163) portion of *Pvalb*+ cells than GCaMP6f (in Ai148) (Fig. 4E,F).

**Figure 4.**
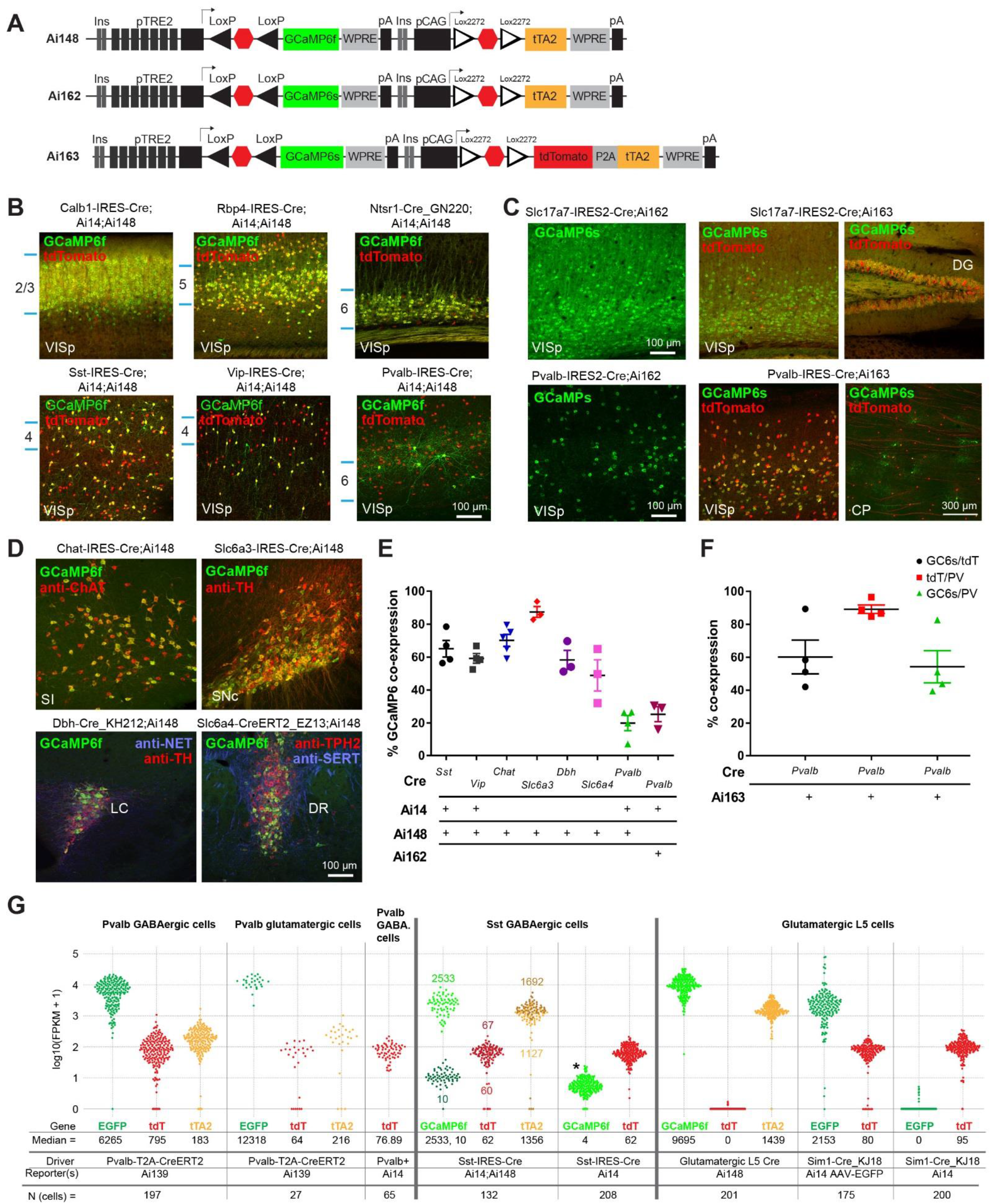
TIGRE2.0 GCaMP6-expressing reporter lines exhibit high-level transgene expression across a broader range of cell types. See also Figures S3-S6. (**A**) Schematic diagram of three new Cre-dependent GCaMP6-expressing TIGRE2.0 reporter lines. (**B**) Representative confocal images of native GCaMP6f and tdTomato fluorescence in indicated Cre;Ai14;Ai148 triple transgenic animals demonstrate layer-specific expression patterns (upper panels) and expression in major inhibitory neuron classes (lower panels). (**C**) Representative confocal images of native GCaMP6s (Ai162) or native GCaMP6s and tdTomato (Ai163) fluorescence in various brain regions of Cre;Ai162 or Cre;Ai163 double transgenic animals. VISp, primary visual cortex. DG, dentate gyrus. CP, caudoputamen. (**D**) Co-localization of native GCaMP6f fluorescence and immunostaining for major neuromodulatory markers in indicated Cre;Ai148 double transgenic animals. In Chat-IRES-Cre;Ai148, cholinergic neurons are labeled by anti-ChAT in substantia innominata (SI). In Slc6a3-IRES-Cre;Ai148, dopaminergic neurons are labeled by anti-TH in substantia nigra, compact part (SNc). In Dbh-Cre_KH212;Ai148, noradrenergic neurons are labeled by anti-NET and anti-TH in locus ceruleus (LC). In Slc6a4-CreERT2_EZ13;Ai148, serotoninergic neurons are labeled by anti-Tph2 and anti-SERT in dorsal raphe (DR). See also Figures S4-S5. (**E**) Quantification of GCaMP6f or GCaMP6s protein expression, in Ai148 or Ai162, respectively, across the indicated Cre-defined cell classes. For cortical interneuron classes (Sst, Vip and Pvalb), native GCaMP6 coexpression with tdTomato (from Ai14) was quantified within VISp and expressed as a percentage of the total number of tdTomato-positive cells. For neuromodulatory neuron types, native GCaMP6f co-expression with the indicated cell-type specific markers in **D** was quantified and expressed as a percentage of the total number of marker-positive cells. Data are presented as mean ± S.E.M; n = 3-5 animals. (**F**) Quantification of GCaMP6s and tdTomato protein co-expression in Pvalb-IRES-Cre;Ai163 mice within VISp. Native GCaMP6s co-expression with tdTomato (black), tdTomato co-expression with anti-PV staining (red), and GCaMP6s co-expression with anti-PV staining (green) was quantified and expressed as percentages. Data are presented as mean ± S.E.M; n = 3-4 animals. (**G**) Quantification of mRNA expression for reporter transgenes by single-cell RNA-seq using the SMART-Seq v4 method on FACS-isolated single cells. Normalized expression values are shown for each cell as fragments per kilobase per million reads (FPKM). Pvalb-T2A-CreERT2;Ai139 cells were separated into two groups by neuronal cell classes (GABAergic vs. glutamatergic). *Pvalb*+;Ai14 cells were collected from pan-GABAergic Cre lines Gad2-IRES-Cre;Ai14 or Slc32a1-IRES-Cre;Ai14 animals, then were selected based on *Pvalb* transcript expression. Glutamatergic L5 Ai148 cells were collected from a Cre line that labels layer 5 excitatory neurons. Sim1-Cre_KJ18;Ai14 AAV-EGFP cells were labeled by injection of an AAV containing pCAG-FLEX-EGFP-WPRE-pA into Sim1-Cre_KJ18;Ai14 mice. Cells were isolated 2 weeks after AAV injection. All cells were FACS-isolated with selection for one of the fluorescent proteins: green fluorescence for Pvalb-T2A-CreERT2;Ai139, Glutamatergic L5 Ai148, and Sim1-Cre_KJ18;Ai14 AAV-EGFP cells; red fluorescence for *Pvalb*+;Ai14, Sst-IRES-Cre;Ai14, and Sim1-Cre_KJ18;Ai14 cells; and either green or red fluorescence for Sst-IRES-Cre;Ai14;Ai148 cells. Because of the green or red fluorescence-based sorting, the Sst-IRES-Cre;Ai14;Ai148 cell population contains two distinct groups of cells, the first with high expression of GCaMP6f (light green, median 2533) and the second with baseline expression (dark green, median 10). These two groups of cells have slightly different, and opposite, levels of tTA2 expression (first group, light brown, median 1127; second group, dark brown, median 1692). Transcriptomic analysis reveals that both groups of cells contain a variety of Sst subtypes (data not shown), and no significant subtype specificity is observed between the two groups, thus the differential expression level of GCaMP6f appears to have occurred randomly. *, a small number of reads from tdTomato expressed by Ai14 are misaligned to the GCaMP6f transcript by the alignment algorithm, thus these are not true GCaMP6f transcripts.

We further investigated Ai148 GCaMP6f expression in subcortical neuromodulatory neurons by quantifying the fraction of cells expressing GCaMP6f driven by corresponding Cre lines in cholinergic neurons in basal forebrain, dopaminergic neurons in ventral tegmental area (VTA) and substantia nigra compacta (SNc), noradrenergic neurons in locus ceruleus (LC), and serotoninergic neurons in dorsal raphe nucleus (DR) identified by immunohistochemistry (Fig. 4D,E and Fig. S4). A large fraction of neuromodulatory neurons displayed strong native GCaMP6f fluorescence that enabled easy cell counting without enhancement by immunostaining (Fig. 4E and Fig. S4). This is a substantial improvement over previous generations of both Rosa26 and TIGRE1.0 GCaMP6 reporter lines, in which GCaMP6 fluorescence was virtually undetectable in neuromodulatory neurons, as shown by the example of the cholinergic neurons labeled by the same Chat-IRES-Cre line (Fig. S5).

To quantitatively evaluate transgene expression levels in TIGRE2.0 lines, we isolated individual cells from different mouse lines by fluorescence-activated cell sorting (FACS) and performed single-cell RNA sequencing using the SMART-Seq v4 method (Tasic et al., 2016) (Fig. 4G). Comparison of GFP and tdTomato mRNA levels in both *Pvalb*+ interneurons and excitatory neurons from Pvalb-T2A-CreERT2;Ai139 mice with those in comparable *Pvalb*+;Ai14 neurons revealed similar tdTomato mRNA levels between the two CAG-driven tdTomato reporters in Ai139 (a TIGRE2.0 line) and Ai14 (a Rosa26 line), but substantially higher (~100 fold) TRE2-driven GFP reporter levels in Ai139. Similarly in layer 5 excitatory neurons, transcript levels of GCaMP6f in Ai148 are ~100 fold higher than those of tdTomato in Ai14. Slightly less but still substantial increase of GCaMP6f transcript levels (~40 fold) compared to tdTomato levels was seen in Sst-IRES-Cre;Ai14;Ai148 and Sst-IRES-Cre;Ai14 animals, although heterogeneity in Sst+ interneurons was also observed. We further found that GFP or GCaMP6f in TIGRE2.0 lines exhibited ~4.5-fold higher expression than GFP from a similar population of layer 5 excitatory neurons infected with a strong, Cre-dependent adeno-associated viral vector AAV-pCAG-FLEX-EGFP-WPRE-pA. These fold changes are higher than what was previously observed between Ai62, a TIGRE1.0 reporter line, and Ai14 (Madisen et al., 2015).

To determine if this high-level transgene expression broadly altered the transcriptomes of Cre+ cells, we examined transcriptome-wide gene expression changes in reporter-positive cells from Pvalb-T2A-CreERT2;Ai139 and Sst-IRES-Cre;Ai14;Ai148 cells, as well as cells from Sim1-Cre_KJ18;Ai14 mice infected with AAV-pCAG-FLEX-EGFP-WPRE-pA. We compared the single cell transcriptomes from these mice to those from matching classes of cells from Cre x Ai14 mice, and found no substantial change in global gene expression and furthermore, fewer differentially expressed genes in TIGRE2.0 cells than in AAV-infected cells (Fig. S6).

We also generated two TIGRE2.0 lines expressing new opsins: Ai167(TIT2L-ChrimsonR-tdT-ICL-tTA2) and Ai168(TIT2L-oChIEF-P2A-tdT-ICL-tTA2) (Fig. 5A). In *Cux2*+ layer 2/3 neurons, strong membrane fluorescence for the ChrimsonR-tdTomato fusion protein was observed (Fig. 5B). In this case, we also show that TIGRE2.0 lines can be combined with TIGRE1.0 lines to achieve dual expression of two reporter genes from a single Cre-dependent tTA2 expression unit, using Cux2-CreERT2;Ai93;Ai167 as an example in which both ChrimsonR-tdT (from Ai167) and GCaMP6f (from Ai93) were expressed in the same or different Cre+ cells. In Ai168, oChIEF, specifically oChIEFAC (Ting et al., 2014), was expressed singly and not as a fusion protein with a fluorescence tag, so its expression was revealed by immunostaining using an antibody against the 2A peptide and it coincided well with the native tdTomato fluorescence (Fig. 5C).

**Figure 5.**
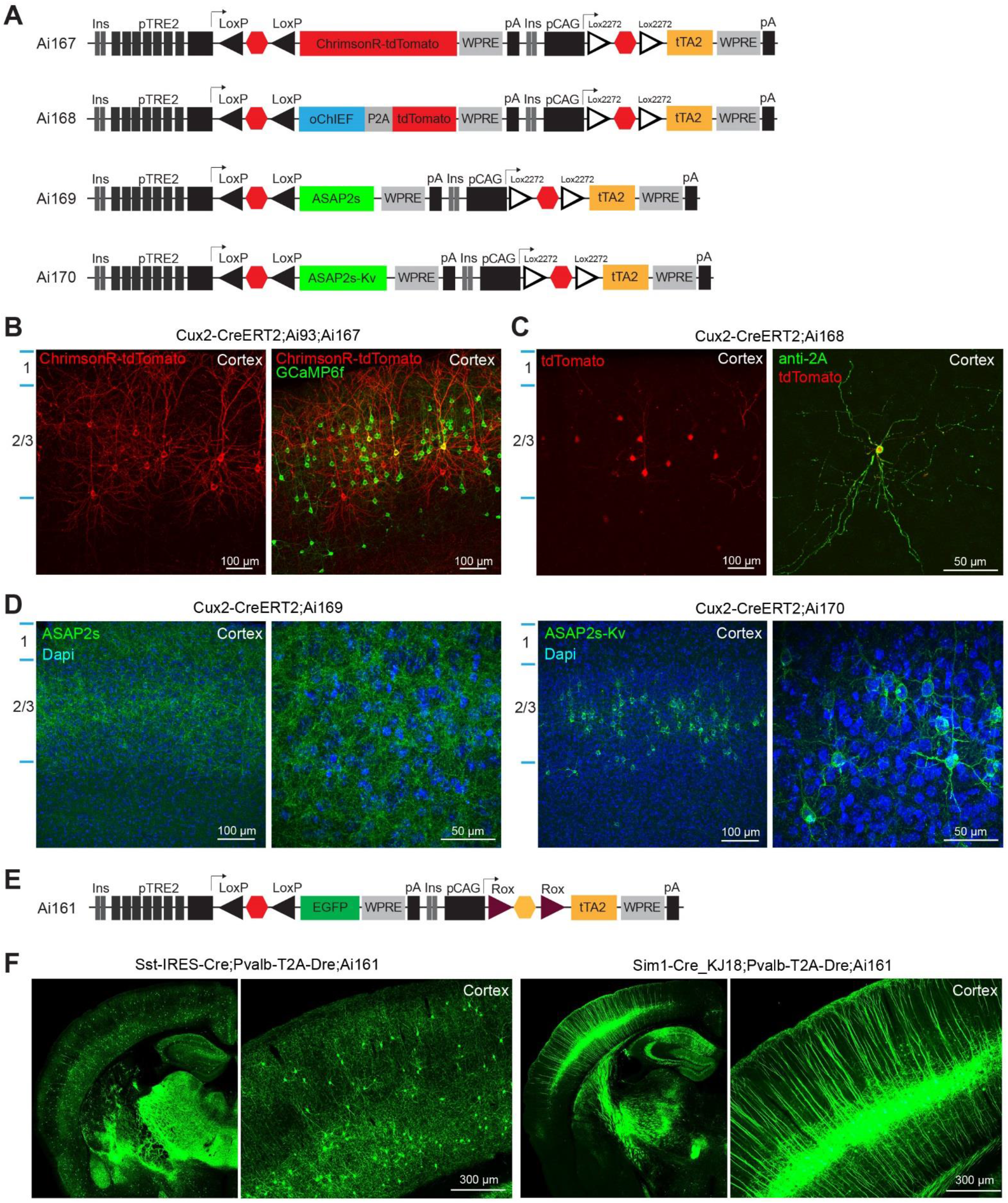
TIGRE2.0 reporter lines with expression of new opsin variants, voltage sensors or intersectional control. (**A**) Schematic diagram of four TIGRE2.0 reporter lines. Ai167 expresses a red opsin variant ChrimsonR fused with tdTomato. Ai168 expresses a blue opsin variant oChIEF along with tdTomato via a P2A sequence. Ai169 expresses a genetically encoded voltage indicator (GEVI) ASAP2s. Ai170 expresses a variant of ASAP2s with a Kv2.1 tag for soma-targeting. See also Figure S7. (**B**) Representative confocal images of native fluorescence show robust expression of ChrimsonR-tdTomato and co-expression with GCaMP6f in a Cux2-CreERT2;Ai93;Ai167 triple transgenic mouse with low-dose tamoxifen induction, indicating tTA2 can activate TRE-promoter driven transgene expression in both cis- and trans-alleles. (**C**) Expression of oChIEF as visualized by anti-2A peptide staining and its co-localization with tdTomato native fluorescence in a Cux2-CreERT2;Ai168 mouse. (**D**) Representative confocal images of native ASAP2s and ASAP2s-Kv fluorescence in Cux2-CreERT2;Ai169 and Cux2-CreERT2;Ai170 mice. As expected, ASAP2s in Ai169 exhibits membrane-localized fluorescence in neuronal processes, and ASAP2s-Kv in Ai170 is enriched in somata. ASAP2s-Kv fluorescence appears brighter than ASAP2s in these mice. (**E-F**) Schematic diagram of a Cre and Dre dependent intersectional TIGRE2.0 reporter line, Ai161 (E), and high-level expression of GFP (imaged by the TissueCyte system) in differential neuronal populations defined by two intersectional crosses (F). In Sst-IRES-Cre;Pvalb-T2A-Dre;Ai161 mice, GFP labels *Sst*+/*Pvalb*+ interneurons, whereas in Sim1-Cre_KJ18;Pvalb-T2A-Dre;Ai161 mice, GFP labels *Pvalb*+ subcortical-projecting layer 5 excitatory neurons.

We also generated two TIGRE2.0 lines expressing a voltage sensor ASAP2s (Chamberland et al., 2017; Yang et al., 2016): Ai169(TIT2L-ASAP2s-ICL-tTA2) expresses ASAP2s, whereas Ai170(TIT2L-ASAP2s-Kv-ICL-tTA2) expresses ASAP2s tagged with a 65-amino-acid cytosolic segment of Kv2.1 to restrict protein localization to the soma and proximal dendrites (Baker et al., 2016; Lim et al., 2000; Wu et al., 2013) (Fig. 5A and Fig. S7). Both lines exhibit robust fluorescence with expected localizations – membrane-localized throughout neuronal processes for ASAP2s in Ai169 and soma-enriched for ASAP2s-Kv in Ai170 (Fig. 5D).

Finally, we generated a Cre and Dre-dependent GFP-expressing intersectional TIGRE2.0 reporter line: Ai161(TIT2L-GFP-ICR-tTA2) (‘R’ stands for RoxStop2Rox), in which tTA2 is under Dre control (Fig. 5E). This transgene expresses GFP in a Cre- and Dre-dependent manner (Fig. 5F). In addition, due to the low-level promiscuous recombination activity of Cre on the Rox sites, in various Cre × Ai161 double transgenic mice, reporter GFP expression was detected in a small number of cells. This promiscuous recombination occurred more frequently in subcortical areas (data not shown).

### Imaging *in vivo* calcium dynamics in cortical inhibitory and excitatory neurons using TIGRE2.0 GCaMP6 reporter mice

Our previously developed GCaMP6-expressing reporter lines (Ai93/94/95/96) displayed strong fluorescence when crossed to various cortical excitatory driver lines (Madisen et al., 2015). However, the GCaMP6 expression from these lines was suboptimal in GABAergic inhibitory interneurons (INs): both the level of GCaMP6 fluorescence and the numbers of active neurons observable were low (Madisen et al., 2015), making *in vivo* functional imaging of INs difficult. Because INs play critical roles in normal brain functions and disorders (Marin, 2012), new reporter lines such as Ai148, Ai162 and Ai163 are urgently needed to effectively study IN functions.

To evaluate the utility of these mice in studying INs, *in vivo* two-photon calcium (Ca^2+^) imaging was conducted in awake, head-fixed, triple transgenic mice containing Ai14 and Ai148 reporters and one of the Cre lines specific for Pvalb, Sst or Vip INs. In all these cases, GCaMP6f was coexpressed with a red, Ca^2+^-insensitive cytosolic fluorescent protein tdTomato (Madisen et al., 2010). Co-expressing tdTomato with GCaMP6 facilitated the localization of and Ca^2+^ imaging in those INs with low neural activity/fluorescence, and also provided a control for the cell-type specificity of GCaMP6f expression in INs (Fig. 4B).

Intracellular Ca^2+^ dynamics was imaged in superficial cortical layers (L) 2/3 (110-350 μm underneath the pia) through an implanted cranial window over the left primary visual cortex (VISp) in head-fixed, awake mice. Consistent with our histological data (Fig. 4B), we observed largely overlapping expression of GCaMP6 with tdTomato across the various imaging depths and Ca^2+^ imaging confirmed that Ai148/162/163 mice greatly improved the expression of GCaMP6 in INs with high specificity. Apparent baseline and/or visually-evoked fluorescence was observed in VISp L2/3 INs (Fig. 6A-C), in sharp contrast to the absent or low expression in corresponding Ai93/Ai95 mice (Madisen et al., 2015). We imaged GCaMP6 responses to fullscreen sinusoidal drifting gratings of 8 orientations (0° to 315° in 45° increments), 3 or 4 spatial frequencies (SFs, [0.02, 0.04, 0.08] or [0.02, 0.04, 0.08 and 0.16] cycle per degree) and 1 temporal frequency (TF, 2 Hz). Overall Ai148/162/163 mice delivered much improved fluorescence signals both in wakefulness (Figs. S8-S10) and under anesthesia (data not shown). Sst-IRES-Cre;Ai14;Ai148 mice (n = 2) showed large (>500%) evoked fluorescence changes (ΔF/F) in response to drifting gratings (Fig. 6A and Fig. S8). We noticed that SST INs exhibited a great variety of baseline and visually evoked fluorescence (Fig. S8), which may reflect the known heterogeneity of SST INs (Urban-Ciecko and Barth, 2016) and/or different levels of neural activity *in vivo*. Similar improvements were found in VIP INs using Vip-IRES-Cre;Ai14;Ai148 mice (n = 3, Fig. 6B and Fig. S9), and the behavioral state (running vs. stationary) dependency of VIP IN activities (Fu et al., 2014) was well depicted in our imaging. In Pvalb-IRES-Cre;Ai14;Ai148 mice, the number of GCaMP6f expressing cells was small (Fig. 4B,E) and Ca^2+^ response was low. Thus we also performed *in vivo* two-photon Ca^2+^ imaging in Pvalb-IRES-Cre;Ai162 and Pvalb-IRES-Cre;Ai163 mice which showed better GCaMP6s expression in PV cells (Fig. 4C,E,F). In these mice, we observed much stronger spontaneous and visually evoked fluorescence responses in PV neurons (n = 1 for Pvalb-IRES-Cre;Ai162, n = 3 for Pvalb-IRES-Cre;Ai163, Fig. 6C and Fig. S10).

**Figure 6.**
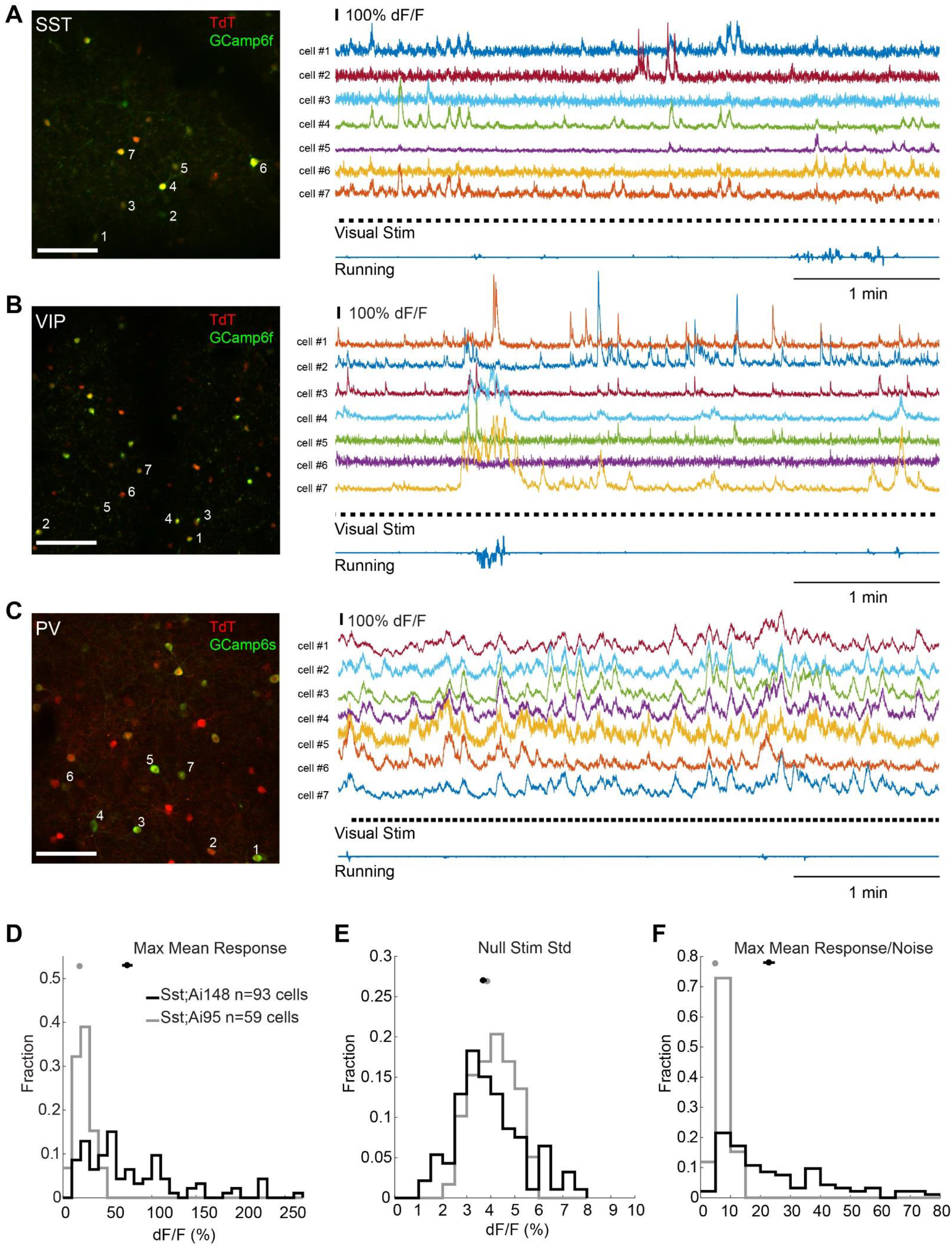
Imaging *in vivo* calcium dynamics in PV, SST and VIP inhibitory interneurons with Ai148 and Ai163 mice. (**A**) Example two-photon Ca imaging of VISp L2/3 SST neurons in an awake, head-fixed Sst-IRES-Cre;Ai14;Ai148 mouse. Left: Z-projection image (time series) showing the entire field-of-view (400 X 400 μm, same for B and C). Imaging was conducted at 212 μm underneath the pial surface. Note the high overlap between tdT and GCaMP6f fluorescence signals. Right: Representative ΔF/F traces for 250 seconds of imaging in 7 SST neurons as labeled in the left panel. Selected neurons covered a wide range of baseline fluorescence. Time course of visual stimuli and running velocity was also shown. Note the response similarity between cells #1, 4 and 7, and response heterogeneity among these 7 cells. (**B**) Example two-photon Ca imaging of VISp L2/3 VIP neurons at 192 μm depth in a Vip-IRES-Cre;Ai14;Ai148 mouse. Left: Z-projection image of the entire field-of-view. Note the nice GCaMP6f expression. Right: Representative ΔF/F traces of 7 VIP cells as labeled in the left panel. Large and sustained responses were observed in cells #4 and 7 during running. Note the fluorescence change started before the initiation of running. (**C**) Example two-photon Ca imaging of VISp L2/3 PV neurons at 280 μm depth in a Pvalb-IRES-Cre;Ai163 mouse. Strong fluorescence responses were observed. The noisier baseline could be due to the high activity of PV neurons *in vivo*. (**D-F**) Improved expression and functionality of GECI in inhibitory interneurons, as shown in a comparison between Sst-IRES-Cre;Ai14;Ai148 (black) and Sst-IRES-Cre;Ai95 (gray) mice. **D**. Population histogram of the mean max responses. **E**. Population histogram of the standard deviation of ΔF/F during gray screens (null stimuli). **F**. Population histogram of the estimated signal-to-noise ratio. Scale bar: 100 μm. Also see Figures S8-S11.

To quantify the improvements of Ai148 compared to Ai95, we compared the fluorescence responses of 93 GCaMP6f/tdT double positive neurons in Sst-IRES-Cre;Ai14;Ai148 mice (n = 2 mice, 8 fields-of-view (FOVs) in VISp) with those of 59 cells in Sst-IRES-Cre;Ai95 mice (n = 2 mice, 6 FOVs in VISp). To control for the imaging noise, we calculated the mean maximum response of individual cells by taking the maximal ΔF/F response averaged within 1 second (0.5 - 1.5 second) post-stimulus onset during the entire recording episode. Compared with Ai95, the distribution of the mean maximum response in Ai148 showed a significant right-ward shift (p < 0.001, Kolmogorov-Smirnov test, Fig. 6D), indicating that a greater portion of Ai148 cells had much stronger fluorescence responses. Consistently, on average the amplitude of Ai148 response was >3-times larger (72.45 ± 5.88% ΔF/F, mean ± SEM, same below) than that in Ai95 (18.93 ± 1.18%, p < 0.001, t-test). In addition, the noise level (3.69 ± 0.15% ΔF/F) in Ai148, which was estimated by the standard deviation of baseline data without visual stimuli, remained comparable with that in Ai95 (3.86 ± 0.11%, p = 0.37, t-test, Fig. 6E). Thus, Ai148 significantly increased the signal/noise ratio by ~4-fold (22.84 ± 1.96 vs. 4.94 ± 0.31, p < 0.001, t-test, Fig. 6F). Strong visual responses were also observed in GCaMP6s-expressing Ai163 and Ai162 lines: in Pvalb-IRES-Cre;Ai163 mice (3 mice, 18 FOVs), for example, responses in PV neurons were on average ~5-fold (52.39 ± 1.05% ΔF/F) greater than that of Pvalb-IRES-Cre;Ai14;Ai148 mice (9.74 ± 1.10%, n = 2 mice, 6 FOVs; p < 0.001, t-test). Although the baseline noise was higher (3.26 ± 0.05% vs. 2.46 ± 0.07%, p < 0.001, t-test) due to the high activity level of PV neurons *in vivo* and increased sensitivity in Ai163, overall the signal/noise ratio significantly increased by ~4-fold compared with Pvalb-IRES-Cre;Ai14;Ai148 (17.42 ± 0.47 vs. 4.48 ± 0.56, p < 0.001, t-test, Fig. S10D).

To further evaluate the functionality of GECIs, we also examined the orientation tuning curves of the fluorescence responses. Tuning curves of GCaMP6f-expressing SST neurons were close to those in pyramidal neurons (Fig. S8), which is consistent with previous studies (Kerlin et al., 2010; Ma et al., 2010). Within the same FOV, diverse Ca^2+^ responses were found in the local population of SST neurons, *i.e.*, neighboring SST neurons may have Ca^2+^ tuning to different orientations and spatial frequencies (Fig. S8). In addition, activity-related Ca^2+^ flashes can also be seen in processes corresponding to dendrites and axons in Ai148 mice (data not shown). For VIP and PV INs, orientation selectivity could be readily extracted from the fluorescence responses in Vip-IRES-Cre;Ai14;Ai148 mice (Fig. S9) and Pvalb-IRES-Cre;Ai163 mice (Fig. S10). Taken together, the results indicate our new GCaMP6-expressing Ai148/162/163 mice offer an excellent combination of greatly improved Ca^2+^ signals with cell-type specificity in 3 major inhibitory interneuron types for *in vivo* functional studies.

To also assess the utility of Ai148 for imaging excitatory neurons, we crossed Ai148 with Cre-lines targeting deep cortical layers. Ntsr1-Cre_GN220;Ai148 mice expressed GCaMP6f strongly and specifically in the layer 6 across the cortex (Fig. S11A) while Ntsr1-Cre_GN220;Camk2a-tTA;Ai93 mice displayed less specificity, with strong expression in some layer 5 cells as well (Fig. S11D). While imaging in VISp layer 6, both sets of mice showed strong visually evoked responses to drifting gratings (Fig. S11C,F). Comparison of the distribution of response amplitudes between Ai93 and Ai148 in the same neuronal populations showed larger changes in fluorescence in Ai93 mice, both when recording in layer 6 (Ntsr1-Cre_GN220 in Fig. S11G, averaged 25 ± 3% ΔF/F in Ai93, n = 183 cells, 12 ± 1% in Ai148, n = 310 cells) and in layer 5 (Tlx3-Cre_PL56 in Fig. S11H, averaged 20 ± 5% ΔF/F in Ai93, n = 281 cells, 12 ± 1% in Ai148, n=349 cells). However, many more GCaMP6f expressing cells were detected overall in both excitatory populations with Ai148, suggesting that this reporter may have detected more cells with no or low evoked activities.

### Light-response properties of TIGRE1.0 and TIGRE2.0 optogenetic mice

We characterized the light-responses of the three new TIGRE1.0 lines and two new TIGRE2.0 lines expressing various channelrhodopsin variants in acute slices of primary visual cortex (VISp). The TIGRE2.0 lines, Ai167 (expressing ChrimsonR-tdTomato) and Ai168 (expressing oChIEF-P2A-tdTomato), were crossed to Cux2-CreERT2, induced with Tamoxifen, and photoresponses from layer 2/3 neurons were measured. We also crossed the TIGRE1.0 lines Ai90 (expressing Chronos-EGFP), Ai134 (expressing ChR2H134R-EYFP) and Ai136 (expressing ReaChR-EYFP) to Scnn1a-Tg3-Cre and ROSA26-ZtTA and measured photo-responses from layer 4 neurons. We stimulated neurons from Ai90, Ai134 and Ai168 mice using a blue LED (470 nm). The reported photo-activation spectra of ReaChR and ChrimsonR are red shifted compared to ChR2 (Klapoetke et al., 2014; Lin et al., 2013); therefore, we utilized a red-orange LED (590 nm) to stimulate cells in the Ai136 and Ai167 mice. Light evoked responses from the Cux2-CreERT2;Ai168 cross, as well as a Pvalb-IRES-Cre;Ai168 cross (both expressing oChIEF), were small and variable (Fig. S12A). Therefore, we did not characterize the Ai168 line further.

We began our characterization of the remaining TIGRE2.0 line and TIGRE1.0 lines by recording photocurrents evoked from opsin-positive neurons. Representative currents from single cells for each reporter line are normalized in amplitude and shown in Fig. 7A. As observed in the initial description of each opsin, the currents carried by these 4 variants demonstrate distinct kinetics (Klapoetke et al., 2014; Lin et al., 2013). Photocurrents from Chronos-positive cells (Ai90) displayed the most rapid kinetics with a sub-millisecond time constant of activation (Fig. S12B). By comparison, activation kinetics were slightly slower in ChrimsonR and ChR2-expressing neurons and dramatically slower in ReaChR-positive neurons. We also observed deactivation kinetics unique to each opsin (Fig. S12C), with the fastest average time constant observed in Chronos, followed by ChR2, ChrimsonR and finally ReaChR. Chronos and ChR2 mediated photocurrents both displayed significant desensitization over the course of a 100 ms stimulus, while desensitization was significantly lower in ChrimsonR and absent in ReaChR photocurrents (Fig. S12D).

**Figure 7.**
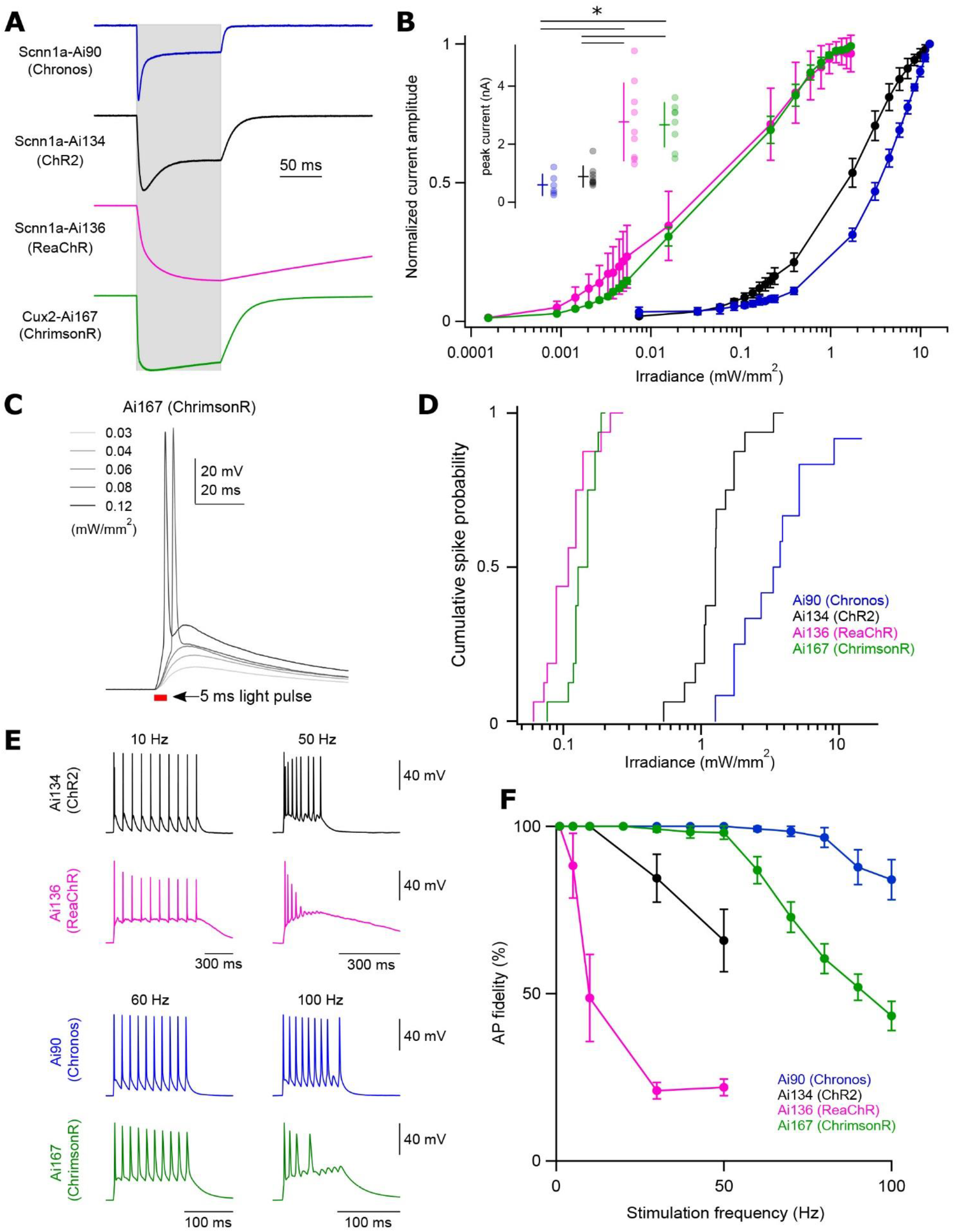
Characterization of light-evoked responses in opsin-expressing TIGRE1.0 and TIGRE2.0 lines. Transgenic mouse strains examined are Scnn1a-Tg3-Cre;ROSA26-ZtTA;Ai90 (simplified as Scnn1a-Ai90 or Ai90, same below) expressing Chronos in cortical layer 4 neurons, Scnn1a-Tg3-Cre;ROSA26-ZtTA;Ai134 expressing ChR2(H134R)-EYFP in cortical layer 4 neurons, Scnn1a-Tg3-Cre;ROSA26-ZtTA;Ai136 expressing ReaChR in cortical layer 4 neurons, and Cux2-CreERT2;Ai167 in cortical layer 2/3/4 neurons. For all data presented here, neurons recorded from Ai90 and Ai134 animals were stimulated with 470 nm light, while experiments characterizing Ai136 and Ai167 mice used 590 nm light. See also Figure S12. (**A**) Representative examples of light-evoked currents in response to 100 ms stimulation (indicated by grey shading). Traces are from individual neurons and are normalized to their peak current amplitude. (**B**) Power-dependence of photocurrent activation for each transgenic line. Data are plotted as mean ± standard deviation (n = 6-9 cells for each line). Color scheme as in **A**. *inset*: Average ± standard deviation of peak current amplitude and data points for individual cells from each transgenic line. Ai90: 0.59 ± 0.36 nA; Ai134: 0.88 ± 0.36 nA; Ai136: 2.76 ± 1.34 nA; Ai167: 2.66 ± 0.76 nA. *P < 0.005. (**C**) Representative responses of a Cux2-CreERT2;Ai167 cell to 5 ms light pulses of increasing power. The timing of the light pulse is indicated by the red bar. (**D**) Cumulative spike probability plotted against photostimulation power for each transgenic line. Significant differences in probability distributions were found between all transgenic lines except Ai136 and Ai167. Kolmogorov-Smirnov Test, *P* < 0.0001 for all comparisons, except Ai136 and Ai167, of which *P* = 0.01. (**E**) Comparison of high frequency firing between transgenic lines. Representative voltage recordings from individual cells in response to 10 repeated stimuli at the indicated frequencies. (**F**) Plot of AP fidelity versus stimulus frequency for each transgenic line. Data are plotted as the mean ± s.e.m (n = 9-15 cells for each line).

The Ai167 TIGRE2.0 line exhibited large-amplitude ChrimsonR-mediated photocurrents (Fig. 7B *inset*: peak current amplitude = 2.66 ± 0.76 nA, n = 8). Among the 3 TIGRE1.0 lines tested, Ai136 peak current amplitudes were comparable to those of Ai167, while Ai134 and Ai90 peak currents were significantly lower. To compare variability of opsin expression within transgenic lines, we measured the coefficient of variation (CV) of the peak current amplitude between cells. Notably, the TIGRE2.0 line Ai167 displayed the smallest CV in peak current amplitude (CV = 0.29; compared to CV = 0.62, 0.42 and 0.48 for Ai90, Ai134 and Ai136 respectively), suggesting the TIGRE2.0 lines can provide robust and consistent expression from cell to cell. We compared photosensitivity by measuring currents across a range of light intensities (Fig.7B). All 4 effector lines displayed strong sensitivity to light intensity with the greatest sensitivity seen in ChrimsonR- and ReaChR-expressing neurons (Ai167 and Ai136).

To compare the capacities of these effector lines to produce optically-evoked suprathreshold responses, we measured light-evoked depolarizations in response to a 5 ms optical stimulus of increasing power (Fig. 7C). From these experiments, we determined the minimum power necessary to generate action potentials (APs) and plotted cumulative spike probability versus light intensity for the 4 transgenic lines (Fig. 7D). Very low powers were required to produce APs in ChrimsonR-positive (Ai167: 0.08 to 0.20 mW/mm^2^, median = 0.15 mW/mm^2^) or ReaChR-positive cells (Ai136: 0.06 to 0.27 mW/mm^2^, median = 0.11 mW/mm^2^). Chronos-positive and ChR2-positive cells required 10 to 30-fold higher powers to evoke APs (Ai90: 1.3 to 14.5 mW/mm^2^; median = 3.5 mW/mm^2^; Ai134: 0.53 to 3.9 mW/mm^2^; median = 1.3 mW/mm^2^). While the required powers are significantly larger, they are readily achievable with commonly utilized light sources, and we observed spiking in all ChR2-positive neurons (n = 16) and all but one Chronos-positive neurons (n = 13, Fig. S12A). ChR2- and Chronos-expressing neurons were excited with a blue (475 nm) LED, slightly shorter than the reported excitation peak of Chronos (500 nm) (Klapoetke et al., 2014). Use of a light source tailored to Chronos would likely provide robust spiking at even lower powers. These data correspond well with our direct recordings of opsin-mediated currents (compare Fig. 7B to 7D). Cells from the Ai136 and Ai167 lines carried the largest and most photo-sensitive currents and produced APs at low light intensities. By comparison, we recorded significantly smaller photocurrents from Ai90 and Ai134 animals and cells from these lines required higher powers to reach AP threshold.

We next utilized light pulses of saturating intensity to determine the minimum duration of light exposure necessary to elicit APs in each effector line (Fig. S12E). ChrimsonR and ReaChR-expressing neurons (Ai167 and Ai136) were highly responsive – all cells exhibited minimum duration of exposure ≤ 0.1 ms. While neurons recorded from Ai90 (Chronos) and Ai134 (ChR2) mice are not as likely to generate APs to such very brief pulses, minimum durations of exposure were below 0.5 ms in nearly all neurons (Fig. S12F). As an additional comparison of optically-evoked spiking across transgenic lines, we presented opsin-expressing neurons with 1 ms light pulses of saturating intensity. We then measured the average latency of AP firing and associated jitter (standard deviation of the latency) associated with this common stimulus. ChrimsonR-positive neurons (Ai167) displayed the lowest latency and jitter (Fig. S12G,H). Notably, under these conditions, all 4 lines displayed highly precise firing with average jitters < 0.1 ms.

Finally, we measured the fidelity of optically-evoked AP firing in response to 10 repeated stimuli of increasing frequency. Representative traces are shown in Fig. 7E. Consistent with the slow kinetics of channel activation and deactivation, ReaChR-expressing cells (Ai136) displayed the least reliable firing in response to high frequency stimuli, with an average fidelity of 22% in response to 50 Hz trains (n = 10 cells). ChR2-expressing cells (Ai134) performed significantly better, displaying an average fidelity of 66% (n = 15 cells; Ai134 compared to Ai136 at 50 Hz: *P* < 0.0005). ChrimsonR and Chronos (Ai90 and Ai167) both exhibited a remarkable ability to follow optical stimuli delivered up to 50 Hz. Therefore, to better differentiate the two lines, we delivered additional stimuli at frequencies ranging from 60 to 100 Hz. At these elevated frequencies, Chronos-expressing cells displayed significantly higher firing fidelity than ChrimsonR-expressing cells (*P* < 0.0001 at 100 Hz). Altogether, our analyses of photocurrents and optically-evoked depolarizations across these transgenic lines suggest they together provide a powerful and complementary toolset for a variety of optogenetic experiments.

### Assessment of adverse effects associated with extremely high-level transgene expression

Very high transgene expression achieved with either viral or transgenic approaches may cause unwanted phenotypes (Han et al., 2012; Potter et al., 2010) and therefore, one should always be cautious and carefully evaluate cell health in each experimental context. While for the vast majority of Cre x TIGRE2.0 crosses examined (>50), the brains and labeled cells appeared healthy, we did observe some adverse effects in selected crosses (Table S1). For example, in double transgenic animals containing Ai148 and Emx1-IRES-Cre (which mediates pan-cortical and pan-hippocampal expression potentially starting as early as embryonic day E9), GCaMP6f fluorescence was extremely bright, the cortex appeared thinner and the hippocampus was smaller (Fig. S13). The mice also had reduced body weight and died prematurely (typically around 1-3 months of age). To investigate if these phenotypes were due to the widespread and developmental expression of GCaMP6f or tTA2, we fed the mice doxycycline-containing chow throughout the pregnancy and animal’s lifetime. This treatment did not ameliorate the adverse phenotypes (data not shown), suggesting that the observed effects were likely due to the widespread developmental expression of tTA2, a potent transcriptional activator. Furthermore, we observed the same phenotypes when Ai140 was crossed with Emx1-IRES-Cre, which expresses a more innocuous molecule (GFP) rather than a calcium buffer (GCaMP6f).

To determine if tTA2 would be better tolerated if expressed later in development, we generated double-transgenic mice containing Ai148, Ai162 or Ai163 and one of pan-cortical Cre lines with later developmental onset, such as Slc17a7-IRES2-Cre, Camk2a-Cre and Camk2a-CreERT2. Encouragingly, double-transgenic animals from the majority of these crosses had no gross behavioral phenotype and the brains appeared largely normal, with the exception of Camk2a-Cre;Ai148 and Slc17a7-IRES2-Cre;Ai148 animals. The cortex from these animals appeared thinner and the dentate gyrus of the latter had notable disorganization (Fig. S13). We also observed an increase in DAPI-negative dark areas, mainly in the hippocampus of some of these double transgenic mice, which may indicate neuronal loss and/or increased gliosis/vasculaturization. Further evaluation of the brains from these crosses using biomarkers for neuroinflammation and quantitative measures to assess brain/subregion volume may help to interpret these more subtle phenotypes or rule them out.

To obtain pan-GABAergic expression, we crossed Ai139, Ai140, and Ai148 reporters with Gad2-IRES-Cre, but never obtained any double positive transgenic mice after genotyping dozens of pups, suggesting embryonic lethality. It is unclear how Gad2-IRES-Cre expression led to lethality, given that double transgenic mice for other interneuron Cre lines (*i.e*., Pvalb, Sst and Vip) all appeared normal. We also extremely rarely obtained double transgenic mice for Ai140 and Sim1-Cre_KJ18, suggesting perinatal lethality; the few double transgenic mice that survived had forelimb atrophy. Sim1-Cre_KJ18 is a BAC transgenic Cre line that, when crossed to Ai14, gives rise to a specific recombination pattern in the adult brain (which is a cumulative representation of all recombination events in the animal’s life) that is restricted to only a few places – cortical layer 5b, pontine gray, nucleus of the lateral olfactory tract, and parts of hypothalamus and medial amygdala. We suspect that the lethality might be due to expression of tTA2 in other, so far unknown body regions. Collectively, these data and our results with the pan-excitatory crosses, suggest that extremely high tTA2 levels are not well tolerated when broadly expressed in CNS, early in development and/or potentially outside of the CNS. We caution users against using Cre lines with widespread and early developmental expression and suggest using the more specific Cre lines in which we have not observed any adverse phenotypes.

We have recently reported the occurrence of aberrant cortical dynamics in some strains of GCaMP expressing mice (Steinmetz et al., 2017). This is an example of more subtle effects with widespread and strong calcium indicator expression, which could be alleviated by doxycycline treatment, and we have developed methods to detect them. These events resemble the synchronized interictal discharges observed in the context of epilepsy, and are characterized by their repetitive discharge (~0.5 Hz), large amplitude (>10%), brief duration (<200 ms in GCaMP6f, <500 ms in GCaMP6s) and a spatial extent that includes motor and somatosensory regions of cortex. Aberrant activity was typically observed in strains with widespread expression of GCaMP across cortex, in particular when GCaMP expression was driven by Emx1-Cre, *e.g*., the Emx1-IRES-Cre;Camk2a-tTA;Ai93 mice. To screen for the incidence of these events in TIGRE2.0 strains we used wide-field calcium imaging through the intact skull. We did not observe aberrant activity from mice expressing GCaMP6s (Ai162) under the control of Slc17a7-IRES2-Cre (Fig. 8A and Fig. S14A, Slc17a7-IRES2-Cre;Ai162, 0/7 mice), but we were able to detect events from the GCaMP6f reporter (Fig. 8B and Fig. S14B, Slc17a7-IRES2-Cre;Ai148, 3/7 mice). For comparison, we also found a low incidence of aberrant activity in Ai93 under the control of Slc17a7-IRES2-Cre (Fig. 8C and Fig. S14C, Slc17a7-IRES2-Cre;Camk2a-tTA;Ai93, 1/5 mice). Events resembled those from the original report (Steinmetz et al., 2017), and our consistent observations in multiple imaging sessions in Emx1-IRES-Cre;Camk2a-tTA;Ai93 mice (5 consecutive positive observations in each of 5 mice, one example shown in Fig. 8D). In all strains where aberrant activities were observed, they were most prominent in the motor and somatosensory cortical regions (Fig. S14), and we found no relationship between these events and the age of the animal (Table S2). Our analysis indicates that mice with GCaMP6s expression rarely exhibited aberrant activity, with only one available example (Fig. 8E) indicating that our analysis is able to detect these events when reported by the slow calcium indicator.

**Figure 8.**
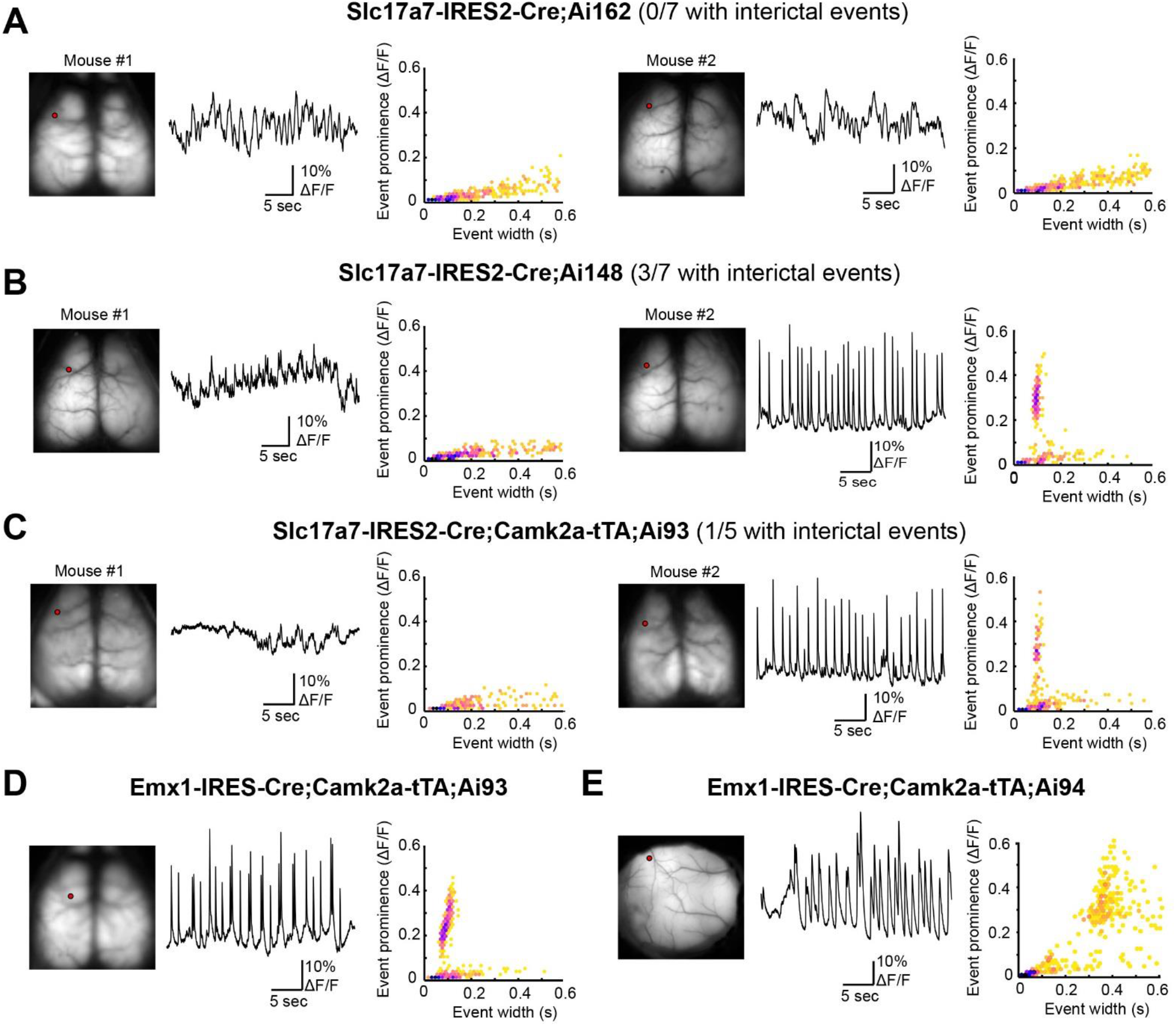
Incidence of aberrant cortical activity in mouse crosses with cortical pan-excitatory GCaMP6 expression. See also Figure S14. (**A**) Wide-field Ca imaging of two example Slc17a7-IRES2-Cre;Ai162 mice. For each mouse, left panel shows GCaMP6 fluorescence from the dorsal surface of the cortex (left), middle panel a 30-second example trace of calcium activity from a location in motor cortex (the point in the left panel), and right panel the relationship between event amplitude (prominence) and event width (hexagonally-binned scatterplot). Same below. None of the 7 mice examined had aberrant activity. (**B**) Wide-field Ca imaging of two example Slc17a7-IRES2-Cre;Ai148 mice. We detected aberrant activity in 3 of 7 mice examined, one of which is mouse #2 shown on the right. (**C**) Wide-field Ca imaging of two example Slc17a7-IRES2-Cre;Camk2a-tTA;Ai93 mice. We detected aberrant activity in 1 of 5 mice examined, and this is mouse #2 shown on the right. (**D**) Wide-field Ca imaging of one example Emx1-IRES-Cre;Camk2a-tTA;Ai93 mouse that had aberrant activity. (**E**) Wide-field Ca imaging of an Emx1-IRES-Cre;Camk2a-tTA;Ai94 mouse. This is the only mouse we examined from this cross that exhibited aberrant activity (region shown is somatosensory cortex). Note the difference in event characteristics when reported by fast (Ai93, D) or slow (Ai94, E) GCaMP6 calcium indicator.

## DISCUSSION

Our previous efforts established the TIGRE locus as a new permissive docking site for insertion of exogenous promoters and transgenes (Madisen et al., 2015). TIGRE1.0 Cre/tTA-dependent intersectional reporter lines demonstrated not only cell type specificity but also much enhanced expression level due to tTA-mediated transcriptional amplification. Currently the availability of tTA driver lines is limited, and in our own several attempts to create new knock-in tTA (specifically, tTA2) driver lines little to no expression was often observed, possibly due to the silencing effect at those endogenous gene loci. Thus we developed the TIGRE2.0 strategy by co-integrating two transgene units, the Cre/tTA-dependent reporter and a CAG promoter-driven, Cre-dependent tTA2 expression unit, into the TIGRE locus. The CAG promoter-driven tTA2 expression in our TIGRE2.0 lines is largely resistant to silencing in nearly all the cell types we have examined, including those we failed to label using TIGRE1.0 lines, such as cortical interneurons and subcortical neuromodulatory neurons, resulting in further enhanced expression of the reporter/effector genes in these cell types that was never reported before. However, the labeling across some of these cell types is still not 100%, indicating the strategy is not perfect yet and silencing may still be an issue as previously reported (Zhu et al., 2007). It will be important to determine in the future if there are specific subtypes that are preferentially unlabeled and to develop strategies to target these particular subtypes.

The TIGRE2.0 platform offers several major improvements over the existing genetic approaches. First, by removing one exogenous driver (tTA), the breeding strategy is simplified to a Cre-dependent approach; it also makes room for the creation of more generally useful dual recombinase-based intersectional strategies such as Cre and Dre reporters (e.g. Ai161) and Cre and Flp reporters (to be generated). Second, as discussed above, by using the robust and ubiquitous CAG promoter to drive tTA2, more consistent and uniform expression of the reporter genes across many different cell types is achieved, minimizing another type of variation. Third, the success of using two different promoters in the TIGRE locus suggests that this locus is indeed universally permissive and it may be used for hosting other promoters or regulatory elements as well.

While the TIGRE2.0 lines represent broadly useful tools for cell type-specific analysis, there are limitations to their utility, as with many existing genetic tools. Among the crosses of TIGRE2.0 reporters with a large number of Cre driver lines, we observed adverse effects with a few specific crosses. The adverse effects observed in several pan-cortical Cre lines were more severe in Cre lines with earlier developmental onset (*e.g*., Emx1-IRES-Cre) and could not be reversed by doxycycline treatment, suggesting that they were caused by the overexpression of tTA2 rather than the effector genes. In general, tTA is a potent transcriptional transactivator that has been reported to cause neurodegeneration when expressed at high levels (Han et al., 2012), thus it is not surprising that its strong expression could be detrimental. In this regard, it is interesting to note that Pvalb-IRES-Cre;Ai163 mice had a higher proportion of PV cells expressing GCaMP6s and also larger *in vivo* GCaMP6s responses than Pvalb-IRES-Cre;Ai162 mice, even though the former (containing tdTomato-P2A-tTA2) likely expresses tTA2 at a lower level than the latter (containing tTA2 alone). This suggests to us that future improvements can be made by moderately reducing tTA2 expression level, and/or by using less potent tTA variants.

Beyond tTA2, widespread and high-level expression of other transgenes could also lead to adverse effects. For example, GCaMP6f mice appear to be more prone to have epileptiform activities than GCaMP6s mice. For the different opsin variants, cells appeared healthier when lower-expressing ROSA26-ZtTA was used compared to Camk2a-tTA to drive Chronos, ChR2 or ReaChR from TIGRE1.0 transgenes. On the other hand, it is interesting to note that ChrimsonR appears to be very well tolerated even in the TIGRE2.0 configuration and exhibits high light sensitivity and high temporal precision. Thus, every molecule likely has its own optimal range of expression and will need to be treated differently to obtain a good balance between maximizing expression/functionality and minimizing toxicity. These conclusions will likely apply to the viral approaches as well. The large set of Rosa26-, TIGRE1.0- and TIGRE2.0-based single-copy transgenic reporter mouse lines we generated has revealed a wide range of expression levels (from insufficient in some cases to “overkill” in others) for a diverse set of molecular tools. They also demonstrate that we now have the ability to fine-tune transgene expression levels as needed. The lessons learned here will be as important as the positive utility demonstrated to facilitate future development and utilization of optimal genetic targeting strategies.

Excellent expression and high functionality has been observed in the majority of the cases where more cell type-specific or more sparsely expressing Cre drivers were used. Our overall results suggest that the diverse set of new driver and reporter transgenic mouse lines reported here can enable numerous cell type-specific applications. These include synaptic terminal labeling (Ai31 and Ai34), nuclear labeling (Ai75 and Ai110), *in vivo* visualization of anatomical structures (Chrm2-tdT), dendritic, axonal and synaptic labeling under electron microscopy (Ai133), full neuronal morphologies (Ai139, Ai140 and Ai161), cortical and subcortical *in vivo* calcium imaging of cell bodies as well as processes (Ai148, Ai162 and Ai163), *in vivo* voltage imaging (Ai86, Ai169 and Ai170), and one- or two-photon optogenetics (Ai40, Ai80, Ai90, Ai134, Ai136 and Ai167). Combined with the large arsenal of Cre, Flp and other driver lines already existing or being generated based on single-cell transcriptome-identified cell types (Ecker et al., 2017; Poulin et al., 2016; Zeng and Sanes, 2017), and furthermore, with complementary viral vectors that allow anterograde or retrograde labeling (Nassi et al., 2015) or systemic delivery (Chan et al., 2017), these applications can be carried out in dense, sparse, intersectional or connectionally-labeled cell populations for very refined circuit dissection, ultimately accelerating our understanding of brain functions in healthy or diseased conditions.

## AUTHOR CONTRIBUTIONS

H.Z., L.M., T.L.D and B.T. designed the transgenic strategies and provided supervision of the project. L.M., T.L.D., H.G. and M.M. generated all new transgenic mouse lines. R.L. and J.H. provided transgenic colony management. J.P. and K.S. managed the genotyping effort. K.S., M.M., K.E.H. and J.A.H. conducted ISH characterization. T.L.D., L.M., H.G., L.S., M.W., E.G. and G.L. conducted histological analysis. R.S.L. and J.W. conducted histological analysis in neuromodulatory mice. B.T., L.T.G., Z.Y. and O.F. conducted single-cell RNA-seq study. N.d.C. and M.M.T. conducted APEX2 staining and electron microscopy study. T.A.H., A.B-M., C.A.B. and G.J.M. conducted patch recording and optogenetic study. M.T.V., D.R.O. and J.W. conducted *in vivo* wide-field calcium imaging. L.L., U.K., L.H. and J.L. conducted *in vivo* two-photon calcium imaging. M.Z.L. and M.C. developed and provided ASAP2s and ASAP2s-Kv constructs. E.S.B. developed and provided Chronos and ChrimsonR constructs. J.T.T. developed and provided oChIEF construct. S.M.S. provided project management and coordination with the Jackson Laboratory. H.Z., T.L.D., L.L., T.A.H., M.T.V., N.d.C., J.L. and B.T. were main contributors to data analysis and manuscript writing, with inputs from other co-authors.

## ACKNOWLEDGMENTS

We are grateful to the In Vivo Sciences, Molecular Biology, Histology, Imaging, and Neurosurgery & Behavior teams at the Allen Institute for their technical support in mouse colony management, genotyping, ISH gene expression characterization, imaging, neurosurgery and behavior training. We thank Robert Hunter for coordinating transgenic mice production at University of Washington. We thank Alice Ting for providing the APEX2 construct, Roger Tsien for providing the tdTomato and ReaChR constructs, Douglas Kim for providing the GCaMP6f and GCaMP6s constructs, Vincent Pieribone for providing the ArcLight construct, Gary Felsenfeld for providing the chicken β-globin HS4 insulator element construct, Philippe Soriano for providing the FlpO construct via Addgene, and Anton Maximov for providing the DHFR-Cre construct. This work was funded by the Allen Institute for Brain Science, NIH grants MH085500 and DA028298 to H.Z., and NIH grant DA036909 to B.T. The authors wish to thank the Allen Institute founder, Paul G. Allen, for his vision, encouragement, and support. The authors declare no conflicts of interest.

## Methods

All experimental procedures related to the use of mice were conducted in accordance with NIH guidelines, and were approved by the Institutional Animal Care and Use Committee (IACUC) of the Allen Institute for Brain Science.

### Transgenic mice generation and transgene expression characterization

All targeting vectors were constructed using gene synthesis and standard molecular cloning approaches. Knock-in driver line and Rosa26-based reporter vectors contained components that were previously described (Madisen et al., 2015; Madisen et al., 2010). For TIGRE1.0- and 2.0-based reporters, vectors containing the following components were constructed: FRT3 - 2X HS4 chicken beta globin insulators – TRE_tight_ (1.0 lines) or TRE2 (2.0 lines) promoter – LoxP – ORF-3X stops – hGH poly(A), PGK polyA) – LoxP – gene – WPRE-bGH poly(A) – 2X HS4 chicken beta globin insulators – CAG promoter – Lox2272 – ORF-3X stops – hGH poly(A), TK poly(A) – Lox 2272 – tTA2 – WPRE – bGH poly(A) – Hygro-SD-FRT5. Targeting of the transgene cassettes into an endogenous gene locus or the Rosa26 locus was accomplished via standard homologous recombination using linearized targeting vector DNA or via CRISPR/Cas9-mediated genome editing using circularized targeting vector in combination with a gene-specific guide vector (Addgene plasmid #42230). Targeting of the transgene cassettes into the TIGRE locus was accomplished via Flp-recombinase mediated cassette exchange (RMCE) using circularized targeting vector and a CAG-FlpE vector (Open Biosystems) as previously described (Madisen et al., 2015). The129S6B6F1 ES cell line, G4 (George et al., 2007), was utilized directly for driver line and Rosa26-based reporter line targeting. A Flp-recombinase landing pad ES cell line derived from G4 cells that was previously described (Madisen et al., 2015) was utilized for RMCE into the TIGRE locus. Correctly targeted ES cells were identified using standard screening approaches (PCR, qPCR, and Southern blots) and injected into blastocysts to obtain chimeras and subsequent germline transmission. Resulting mice were crossed to the Rosa26-PhiC31 mice (JAX Stock # 007743) to delete the pPGK-neo or pPGK-hygro selection marker cassette, and then backcrossed to C57BL/6J mice and maintained in C57BL/6J congenic background.

Induction of DHFR-domain containing dCre, dgCre or dgFlpO driver lines was done by administration via oral gavage (PO) of trimethoprim (TMP) at 0.3 mg/g body weight per day for 3 days. TMP working solution was diluted from a 10x stock in 100% DMSO with a 2% methylcellulose solution. Induction of CreERT2 driver lines was done by administration via oral gavage (PO) of tamoxifen (50 mg/ml in corn oil) at 0.2 mg/g body weight per day for 1-5 days. Mice can be used for experiments starting at 1-2 weeks after TMP or tamoxifen dosing.

Expression of the reporter transgenes was assessed by laser-scanning confocal microscopy and RNA *in situ* hybridization (ISH). All ISH was performed via a standardized production platform that was previously described (Harris et al., 2014; Madisen et al., 2010). Data can be found in the AIBS Transgenic Characterization database (http://connectivity.brain-map.org/transgenic/search/basic).

For parvalbumin or 2A antibody staining, free floating brain sections were rinsed three times in phosphate-buffered saline (PBS), blocked for 1 hour in PBS containing 5% donor donkey serum, 2% bovine serum albumin (BSA) and 0.1% Triton X-100, and incubated overnight at 4°C in the anti-parvalbumin primary antibody (PV, 1:3000, Swant PV235) or in the anti-2A peptide primary antibody (2A, 1:500, Millipore ABS31). The following day, sections were washed three times in PBS, followed by incubation in blocking solution containing an Alexa 647 conjugated secondary antibody (1:250, Jackson 715-605-151), washed in PBS, and mounted in Vectashield containing DAPI (H-1500, Vector Labs). For all other antibody staining, sections were subjected to antigen retrieval with 10 mM sodium citrate pH 6.0 with 0.05% Triton-X-100 (heated to 75°C for 20 minutes), rinsed, and then blocked with 5% normal donkey serum and 0.2% Triton X-100 in PBS for one hour. Tissue was then incubated in primary antibodies diluted in the blocking solution for 48-72 hours at 4°C, washed the following day in 0.2% Triton X-100 in PBS and then incubated in the appropriate Alexa-conjugated secondary antibodies (1:500, ThermoFisher). Sections were then rinsed in 0.2% Triton X-100 in PBS, followed by PBS alone, mounted on gelatin-coated slides and cover-slipped with prolong diamond antifade mounting media (P36965, ThermoFisher). The following primary antibodies were used to detect transgene expression in neuromodulatory cell types: anti-norepinephrine transporter (NET, 1:500, Atlas Antibodies AMAb91116), anti-tyrosine hydroxylase (TH, 1:1000, Abcam ab112), anti-serotonin transporter (SERT, 1:500, EMDMillipore MAB1564), anti-tryptophan hydroxylase 2 (TPH2, 1:250, EMDMillipore ABN60), anti-choline acetyltransferase (ChAT, 1:300, EMDMillipore AB144P). In a subset of experiments, we compared antibody-enhancement to native GCaMP6 fluorescence using an anti-GFP primary antibody (1:5000, Abcam, Ab13970) and donkey anti-chicken secondary antibody (703-545-155, Jackson ImmunoResearch). Images (1024 X 1024) of native or antibody-enhanced fluorescence were collected on an Olympus Fluoview FV1000 confocal microscope or a Leica SP8 resonant-scanning confocal microscope and were manually counted for each experimental condition using the cell counter plugin within ImageJ (K. De Vos, University of Sheffield) by observers blind to the genotype of the animals. Data were analyzed and graphed using GraphPrism 7.02 software (Hearne Scientific Software, Chicago, IL).

### Quantification of reporter transgene expression by single-cell RNA-sequencing

For comparison of reporter transgene expression level, two Sim1-Cre_KJ18;Ai14 mice, one male and one female, at P43, were injected with 200 nl of adeno-associated virus AAV-pCAG-FLEX-EGFP-WPRE-pA into the anterolateral motor cortex (ALM) by stereotaxic injection in both hemispheres at coordinates A/P 2.50, M/L 1 and −1, D/V 0.60 using a pressure injection system (Nanoject II, Drummond Scientific Company, Catalog# 3-000-204). AAV-treated cells were collected 14 or 16 days post-infection at P57 or P59.

Cells from transgenic mice or transgenic mice injected with AAV-pCAG-FLEX-EGFP-WPRE-pA were collected by microdissection, single-cell suspension, and single-cell FACS. Cells were then frozen at −80°C, and were later processed for scRNA-seq using the SMART-Seq v4 method, as described in the Allen Institute Transcriptomics Technical Whitepaper (http://help.brain-map.org/download/attachments/8323525/CellTypes_Transcriptomics_Overview.pdf) (Tasic et al., 2016). After sequencing, raw data was quantified using STAR v2.5.3 and were aligned to both a Ref-Seq transcriptome index for the mm10 genome, and a custom index consisting of transgene sequences. FPKM values were calculated by dividing raw counts for each transgene by the length of each gene in kilobases and by the total number of reads that mapped to both the transcripome index and to the transgene sequences in millions of reads.

To examine transcriptome-wide gene expression changes, scRNA-seq datasets were mapped to reference transcriptomic cell types using nearest centroid classifiers. Differentially expressed genes between groups of cells were detected using limma voom mode (Ritchie et al., 2015).

### *In vivo* two-photon calcium imaging in cortical interneurons

Detailed surgical and imaging procedures have been published previously (Madisen et al., 2015). Briefly, adult mice (2 – 5 month old, both genders, n = 9, namely, 2 Sst-IRES-Cre;Ai14;Ai95, 2 Sst-IRES-Cre;Ai14;Ai148, 2 Pvalb-IRES-Cre;Ai14;Ai148 and 3 Vip-IRES-Cre;Ai14;Ai148 mice) were implanted with a metal head-post and a ~5-mm cranial window over the left visual cortical area under anesthesia. After 7 days of recovery from surgery, mice were habituated to head fixation and presentation of visual stimulations on an in-house made running device (a freely-rotating running disc) for 2 weeks before imaging experiment started. During imaging, mice were awake, head-fixed but allowed to run or rest on the freely-rotating running disc with or without visual stimuli presented on a calibrated LCD monitor spanning 60° in elevation and 130° in azimuth to the contralateral (right) eye. For visually evoked responses, whole-screen sinusoidal drifting gratings were presented showing 8 orientations (0° to 315° in 45° increment), 3 or 4 spatial frequencies (SFs, [0.02, 0.04, 0.08] or [0.02, 0.04, 0.08 and 0.16] cycle per degree) and 1 temporal frequency (2 Hz), presented at 80% contrast in a random sequence for 8 repetitions. Each drifting grating lasted for 2 seconds with an interstimulus-interval of 1 or 2 seconds. A gray screen at mean illuminance was presented randomly for 16 times. The mouse’s eye was positioned ~22 cm away from the center of the monitor. Due to the choice of whole-screen visual stimuli, we did not track the eye position.

Two-photon calcium imaging was performed using a Bruker (Prairie) two-photon microscope with 8 kHz resonant scanners, coupled with a Chameleon Ultra II Ti:sapphire laser system (Coherent). In the current study, we crossed the Ai148 reporter lines with Pvalb-, Sst- and Vip-IRES-Cre;Ai14 mice to co-express the GECI GCaMP6f together with a red, calcium insensitive cytosol marker tdTomato in PV, SST and VIP positive inhibitory interneurons, respectively. Coexpression of tdTomato facilitated locating and calcium imaging those INs with low expression of GECI. Fluorescence excited at 920 nm wavelength with <70 mW laser power measured after objective was collected in two spectral channels using green (510/42 nm) and red (641/75 nm) emission filters. The emission filter set we selected ensured good separation between red and green channels. Fluorescence images were acquired at various frame rates (512X512 pixels, 30 Hz; 256X256 pixels, 60 Hz) through a 16x water-immersion objective lens (Nikon, NA 0.8), with or without visual stimulations.

Acquired image data were analyzed with in-house developed Matlab scripts. In summary, data were corrected for in-plane motion artifacts using cross-correlation motion correction method between frames (Dombeck et al., 2007). Image segmentation was done using Independent Component Analysis (ICA) followed by automated Region of Interest (ROI) selection with manual confirmation/correction (Mukamel et al., 2009). Green fluorescence signal of all pixels within each ROI was averaged by image frames. Fluorescence signal from the red channel was not quantitatively analyzed. No neuropil signal subtraction was done for these data. The baseline for ΔF/F calculation was determined from the mode of the F trace.

### *In vivo* two-photon calcium imaging in cortical excitatory neurons

Imaging of the Ai93 and Ai148 reporter lines crossed with Ntsr1-Cre_GN220 and Tlx3-Cre_PL56, which express GCaMP6f in deep cortical layer 6 and 5 respectively, was conducted in the Allen Brain Observatory pipeline. Briefly, adult mice (from P37 to P63, both sexes, n = 7) were implanted with a metal head-post and a ~5-mm cranial window over the left visual cortical area under anesthesia. After 7 days of recovery from surgery, the visual cortex was mapped using intrinsic imaging as described previously (Garrett et al., 2014). An automated segmentation and annotation module was developed to delineate boundaries between visual areas. To provide a reliable targeting of two-photon calcium imaging experiments, a consistent anatomical coordinate corresponding to the center of V1 (which maps to center of the retina) was used.

Mice were then habituated to head fixation and presentation of visual stimulations on an in-house made running device (a freely-rotating running disc) for 2 weeks before imaging experiment started. During imaging, mice were awake, head-fixed but allowed to run or rest on the freely-rotating running disc with or without visual stimuli presented on a calibrated LCD monitor.

The screen spanned 120° X 95° of visual space and was positioned 15 cm from the mouse (118.6 mm lateral, 86.2 mm anterior and 31.6 mm dorsal to the right eye). Each screen (ASUS PA248Q LCD monitor) was gamma calibrated using a USB-650 Red Tide Spectrometer (Ocean Optics). Luminance was measured using a SpectroCAL MKII Spectro radiometer (Cambridge Research Systems). Monitors brightness (30%) and contrast (50%) corresponded to a mean luminance of 50 cd/m2.

For quantifying grating responses, the stimulus consisted of a full field drifting sinusoidal grating at a single spatial frequency (0.04 cycles/degree) and contrast (80%). The grating was presented at 8 different directions (separated by 45°) and at 5 temporal frequencies (1, 2, 4, 8, 15 Hz). Each grating was presented for 2 seconds, followed by 1 second of mean luminance gray before the next grating. Each grating condition (direction & temporal frequency combination) was presented 15 times, in a random order. There were blank sweeps (i.e. mean luminance gray instead of grating) presented approximately once every 20 gratings.

Two-photon calcium imaging was performed using a Nikon A1R MP+. The Nikon system was adapted to provide space to accommodate the behavior apparatus. Laser excitation was provided by a Ti:Sapphire laser (Chameleon Vision – Coherent) at 910 nm. Pre-compensation was set at ~10,000 fs2. Movies were recorded at 30Hz using resonant scanners over a 400-μm field of view. Two-photon movies (512x512 pixels, 30Hz), eye tracking (30Hz), and behavior (30Hz) were recorded and continuously monitored. We imaged the deep cortical layers (layer 5 and 6) through a 16x water-immersion objective lens (Nikon, NA 0.8) using less than 240 mW (measured after objective) to avoid brain damage.

Acquired image data were analyzed with in-house developed scripts. A comprehensive description of the data analysis pipeline is available on the Allen Brain Observatory portal (http://help.brain-map.org/display/observatory/Documentation). Briefly, each pixel of the calcium movie was first corrected for pixel leakage, then each frame was motion corrected to a reference frame. A rule-based segmentation algorithm automatically extracted all ROIs covering active somatic compartments. For each ROI, neuropil signals were then subtracted and nearby ROIs were unmixed. Each ROI associated trace was then converted to ΔF/F. For drifting grating quantification, tuning was calculated from the averaged response over the duration of stimulus presentation.

### Acute slice physiology and optogenetics

Para-sagittal slices were prepared from 6-16 weeks old mice of either sex. Mice were anaesthetized with isoflurane, decapitated and brains were swiftly removed. 350 μm slices were cut using a Leica VT1200S vibratome in ice cold slicing solution. Slicing solution contained 87 mM NMDG, 2.5 mM KCl, 1.2 mM NaH_2_PO_4_, 30 mM NaHCO_3_, 20 mM HEPES, 25 mM glucose, 12 mM N-acetyl cysteine, 5 mM Na-ascorbate, 2 mM thiourea, 3 mM Na-pyruvate, 10 mM MgSO_4_, 0.5 mM CaCl_2_. pH was adjusted to 7.3-7.4 using HCl. Slices were stored in slicing solution for 15 minutes at 34° C, after which they were transferred to artificial cerebrospinal fluid (ACSF) and stored at room temperature until use. ACSF contained 124 mM NaCl, 2.5 mM KCl, 1.2 mM NaH_2_PO_4_, 24 mM NaHCO_3_, 5 mM HEPES, 12.5 mM glucose, 2 mM MgSO_4_ and 2 mM CaCl_2_. pH was adjusted to 7.3-7.4 mM using NaOH, osmolarity = 290-300 mOsm. Prior to use, slices were transferred to a recording chamber continuously perfused with ACSF heated to 32 ± 2° C. Recordings were made using 3 – 5 MΩ borosilicate glass pipettes. Internal solution contained 130 mM K-gluconate, 4 mM KCl, 10 mM HEPES, 0.3 mM EGTA, 10 mM phosphocreatine disodium salt, 4 mM MgATP and 0.3 mM Na-GTP. During recording, 10 μM NBQX was added to ACSF to block excitatory synaptic transmission. Measurements of photocurrents were performed with neurons clamped to −70 mV. 100 nM tetrodotoxin added to the bath to block voltage-gated sodium channels in voltage clamp recordings. Access resistance was <20 MΩ at the start of all experiments and was monitored throughout the experiment. Recordings were terminated if access resistance increased to >25 MΩ. Data were recorded using Multiclamp 700B amplifiers (Axon Instruments) and digitized using ITC-18 data acquisition interfaces (Heka). Data were acquired using custom scripts in Igor Pro (Wavemetrics).

Photostimulation of cells was performed using either a 470 nm or 590 nm LED reflected into a 63x/1.0 NA water immersion objective. The power of the light stimulus was controlled via pulse-width modulation of TTL signals passed to an analog LED driver (LightSpeed Technologies). Powers were measured using a power meter (Thor Labs PM-100A) placed after the objective. Experiments measuring action potential (AP) entrainment at different frequencies utilized different pulse widths for each transgenic line. This was to accommodate the differences in minimum light duration necessary to evoke a single AP (Fig. S9F). Pulse durations utilized were Ai90: 1 ms; Ai134: 0.5 ms, Ai136: 0.05 ms, and Ai167: 0.1 ms. Experiments performed using longer pulses in Ai136 and Ai167 mice generally resulted in multiple APs in response to the first pulse, followed by decreased fidelity at the end of the pulse train.

Analysis was performed using Igor Pro and custom Python (2.7) scripts combined with the Allen Institute Software Development Kit to measure the timing of APs in whole cell recordings. Statistical comparisons of population averages of transgenic lines were made by first performing a one-way ANOVA, followed by an unpaired t-tests between groups with a Bonferroni correction for multiple comparisons (6 comparisons between 4 groups. α = 0.0083).

### Wide-field calcium imaging

Wide-field calcium imaging was performed either through the intact skull using a modification of the method from (Silasi et al., 2016) or through a 5-mm-diameter chronically implanted cranial window centered over the left visual cortex (Andermann et al., 2010; Zhuang et al., 2017). For through-skull imaging, the skull was exposed and cleared of periosteum, and a #1.5 borosilicate coverslip (Electron Microscopy Sciences, #72204-01) was fixed to the skull surface by a layer of clear cement (C&B Metabond, Sun Medical Co.). A 3D-printed light shield was fixed around the coverslip using additional Metabond, and the outward-facing surfaces were coated with an opaque resin (Lang Dental Jetliquid, MediMark). Mice with chronically implanted windows received a 5-mm-diameter craniotomy over the left hemisphere, centered at 2.7 mm lateral and 1.3 mm anterior to lambda. The craniotomy was sealed with a stack of three #1 coverslips, attached to each other using optical adhesive (Norland) and to the skull with Metabond. The window provided optical access to the left visual cortex, the posterior aspect of somatosensory cortex and medial aspect of dorsal retrosplenial cortex. In both cases, a custom-manufactured titanium headpost was fixed posterior to the lightshield/coverslip and dorsal to the cerebellum using Metabond. All surgical procedures were performed under isoflurane anesthesia. Image acquisition used a Hamamatsu Flash4.0 v2 or v3 sCMOS camera running with a 10-ms rolling exposure (100 Hz). Images were produced by a tandem-lens macroscope (Scimedia THT-FLSP) with 1.0x tube and objective lenses (Leica 10450028) for through skull imaging or a 1.0x tube lens paired to a 1.6x objective lens (Leica 10450029) for imaging through the chronically implanted window. Epifluorescence illumination used a 470-nm LED (Thorlabs M470L3) filtered (Semrock FF01-474/27-50) and reflected by a dichroic mirror (Semrock FF495-Di03-50 x70) through the objective lens. Fluorescence emission passed through a filter (Semrock FF01-525/45-50) to the camera. Data were spatially downsampled by averaging, and calcium traces were obtained by subtracting and then dividing each pixel by its mean over the whole time series. At all locations, mice were either male or female and were 7–30 weeks of age.

Traces were analyzed by finding peaks and computing two parameters: prominence and full-width at half-prominence. Prominence is defined as the height of the peak relative to the greater of the two minima between the peak and its surrounding two higher peaks (or beginning/end of trace if no higher peak). Peak detection used PeakUtils 1.0.3 (Python 2.7).

## Supplementary Figures

**Figure S1.**
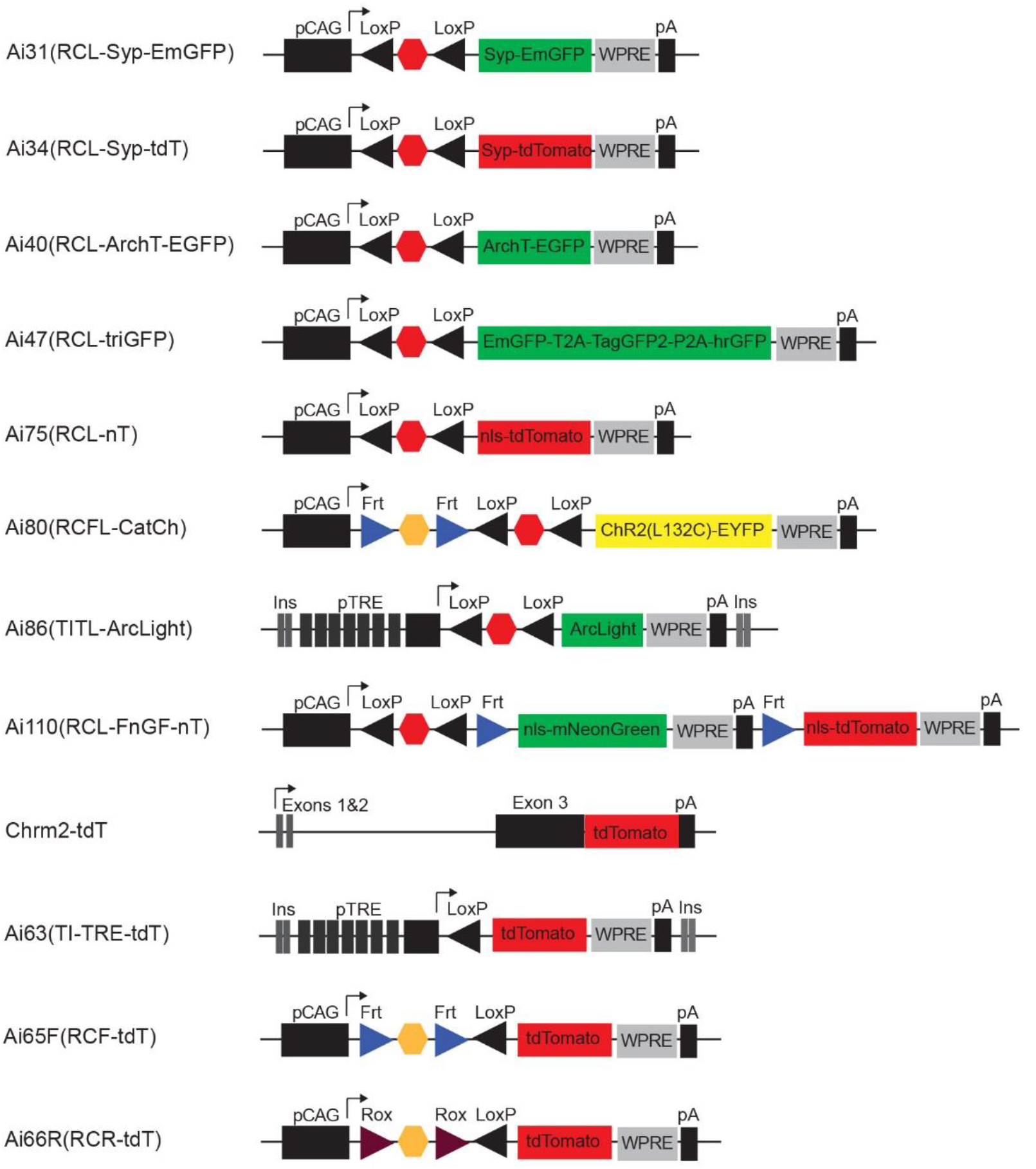
Schematic diagrams of new transgenic reporter lines. Related to Table 2. RCL, Rosa26 – CAG promoter – LoxP-STOP-LoxP. RCFL, Rosa26 – CAG promoter – FRT-STOP-FRT – LoxP-STOP-LoxP. RCF, Rosa26 – CAG promoter – FRT-STOP-FRT. RCR, Rosa26 – CAG promoter – Rox-STOP-Rox. TITL, TIGRE – Insulators – TRE promoter – LoxP-STOP-LoxP. EmGFP, Emerald Green Fluorescent Protein. FnGF, FRT – nls-mNeonGreen – FRT. hrGFP, humanized Renilla Green Fluorescent Protein. nT, nls-tdTomato. Syp, Synaptophysin. tdT, tdTomato.

**Figure S2.**
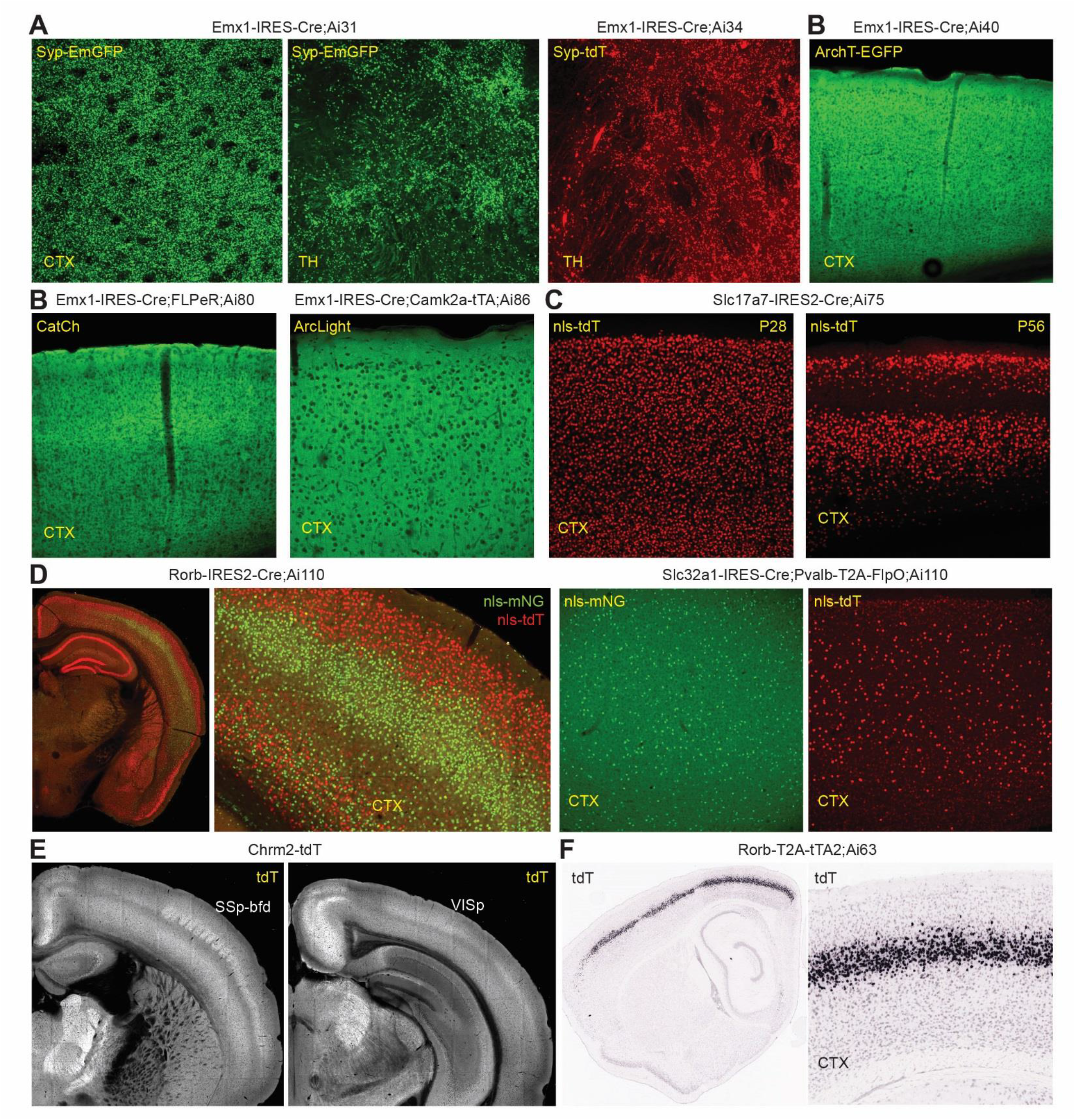
Expression validation of new transgenic reporter lines. Related to Table 2. (**A**) When crossed to the pan-cortical Emx1-IRES-Cre, Ai31 and Ai34 profusely label synaptic terminals by Syp-EmGFP and Syp-tdT, respectively. Native fluorescence was imaged by confocal microcopy in cortex (CTX) and thalamus (TH). (**B**) When crossed to Emx1-IRES-Cre, Ai40, Ai80 and Ai86 display pan-cortical native fluorescence for ArchT, CatCh and ArcLight, respectively. (**C**) When crossed to the glutamatergic Slc17a7-IRES2-Cre, nls-tdTomato from Ai75 labels extremely brightly all cortical glutamatergic nuclei at age P28. However, by P56 nls-tdTomato expression decreases or appears absent in selected cell populations, and continues to decrease in more cells with time. Similar observations have been made when Ai75 was crossed to other Cre lines. The cause of this phenomenon is unknown. Thus we recommend only short-term usage of the Cre-dependent nuclear labeling by Ai75. (**D**) Ai110 was designed as a dual Cre/Flp-dependent reporter to express nls-mNeonGreen (nls-mNG) in cells where only Cre is present, and express nls-tdTomato (nls-tdT) in cells where both Cre and Flp are present. However, there is substantial leaky expression of nls-tdT in the absence of Flp, as seen in Rorb-IRES2-Cre;Ai110 native fluorescence images. The cause of this phenomenon is unknown. On the other hand, the Cre-dependent nls-mNG expression works as expected, as seen in both Rorb-IRES2-Cre;Ai110 and Slc32a1-IRES-Cre;Pvalb-T2A-FlpO;Ai110 native fluorescence images for predominantly layer 4 labeling (Rorb) and pan-GABAergic neuronal labeling (Slc32a1), respectively. The nls-tdT portion can still be useful, as the presence of Flp can turn nls-tdT expression fully on, and this full expression level is clearly distinguishable from the leaky expression (see *Pvalb*-specific nls-tdT labeling in Slc32a1-IRES-Cre;Pvalb-T2A-FlpO;Ai110 in the rightmost panel). (**E**) The knock-in Chrm2-tdT fusion protein provides native fluorescence for anatomical boundaries for certain cortical areas such as barrel cortex (SSp-bfd) and primary visual cortex (VISp), visible *in vivo*. (**F**) RNA *in situ* hybridization (ISH) with tdT probe shows the cortical layer 4 specificity in a Rorb-T2A-tTA2;Ai63 mouse, in which Ai63 is a tTA-dependent reporter (derived from the Cre/tTA dependent TIGRE1.0 reporter Ai62) expressing tdT from the TRE-tight promoter.

**Figure S3.**
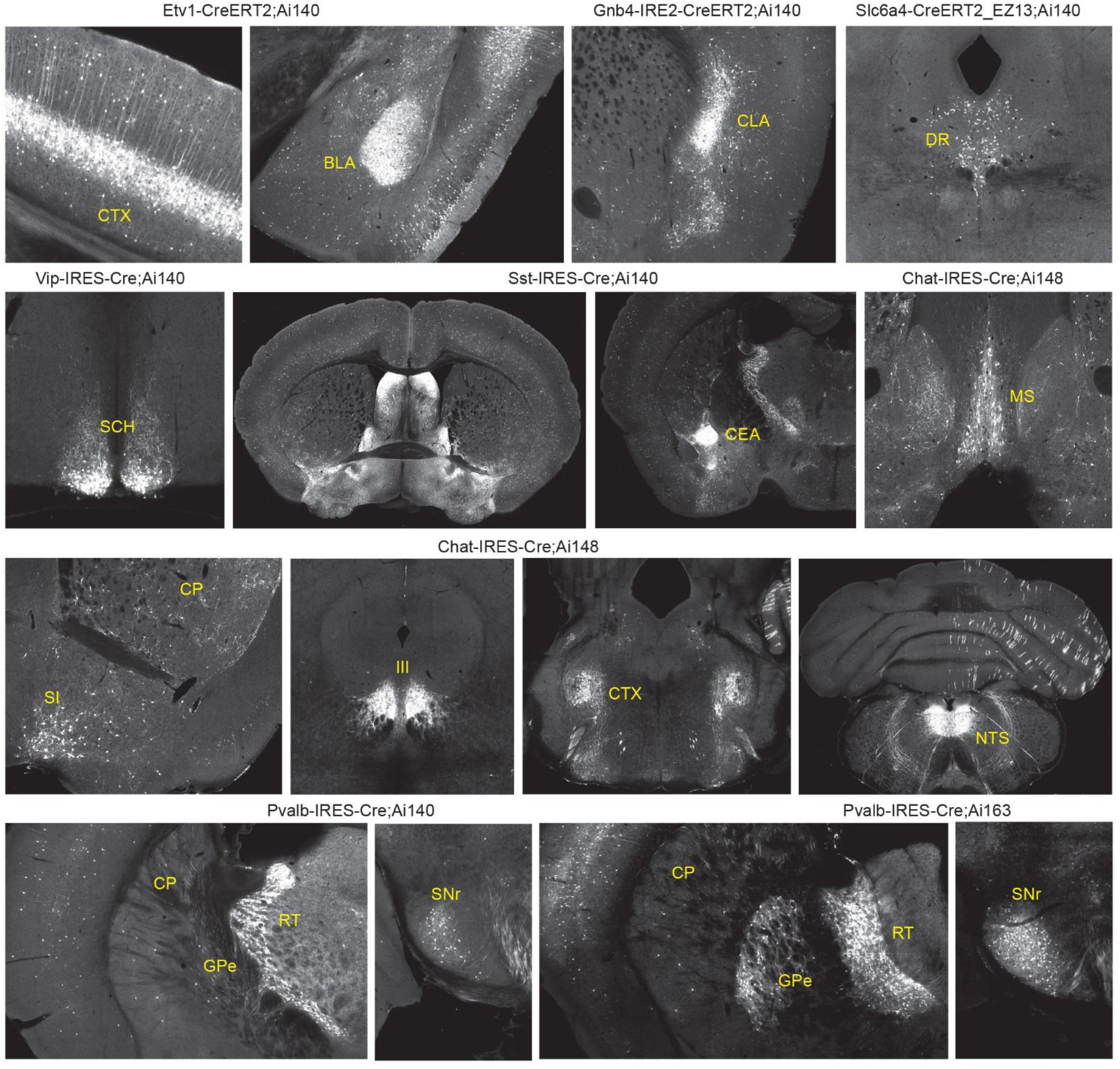
TIGRE2.0 reporter lines exhibit high-level transgene expression across a wide range of cortical and subcortical regions when crossed to a diverse set of Cre driver lines. Related to Figures 3 and 4. EGFP (in Ai140), GCaMP6f (in Ai148) and GCaMP6s (in Ai163) native fluorescence was imaged by the TissueCyte serial two-photon tomography (STPT) system. Note that more cells are labeled in Pvalb-IRES-Cre;Ai163 than in Pvalb-IRES-Cre;Ai140. BLA, basolateral amygdalar nucleus. CEA, central amygdalar nucleus. CLA, claustrum. CP, caudoputamen. CTX, cortex. DR, dorsal nucleus raphe. GPe, globus pallidus, external segment. III, oculomotor nucleus. MS, medial septal nucleus. NTS, nucleus of the solitary tract. RT, reticular nucleus of the thalamus. SCH, suprachiasmatic nucleus. SI, substantia innominata. SNr, substantia nigra, reticular part.

**Figure S4.**
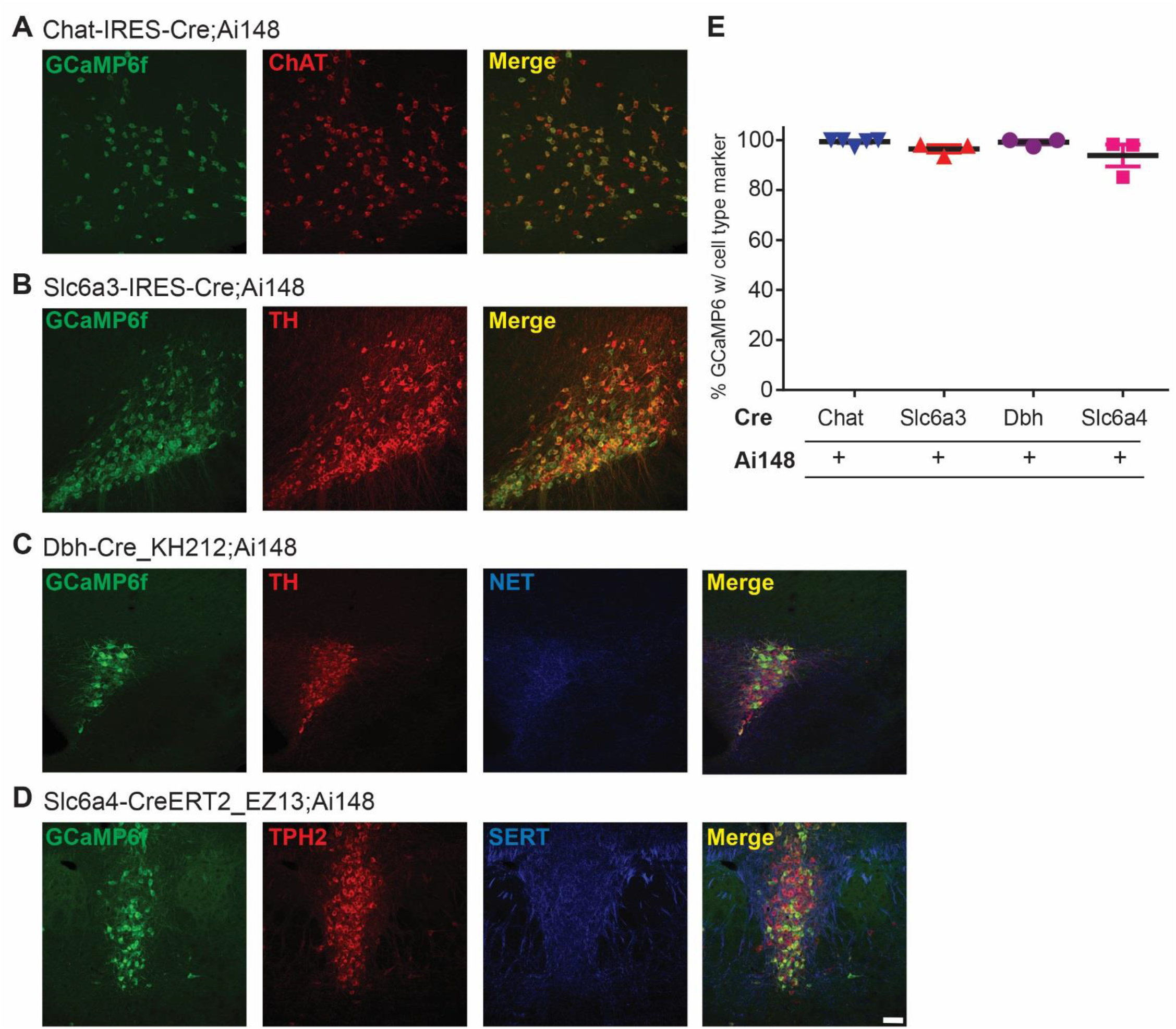
TIGRE2.0 GCaMP6f-expressing reporter line labels neuromodulatory cell types with high specificity. Related to Figure 4. (**A-D**) Representative confocal images show that Ai148 drives specific expression in a variety of neuromodulatory cell types as indicated by co-expression of GCaMP6f with the neuromodulatory cell-type specific markers. The Ai148 reporter line specifically labels cholinergic, choline acetyltransferase (ChAT)-positive neurons in Chat-IRES-Cre crosses (**A**), dopaminergic, tyrosine hydroxylase (TH)-positive neurons in Slc6a3-IRES-Cre crosses (**B**), noradrenergic, TH- and norepinephrine transporter (NET)-positive neurons in Dβh-Cre_KH212 crosses (**C**), and serotonergic, tryptophan hydroxylase 2 (TPH2)- and serotonin transporter (SERT)-positive neurons in Slc6a4-CreERT2_EZ13 crosses (**D**). Slc6a4-CreERT2_EZ13;Ai148 mice received 1-day tamoxifen induction after P20 and survived for at least 2 weeks before perfusion. Scale bar: 50 μm. (**E**) The TIGRE2.0 reporter line Ai148 labels a subset of cells within each neuromodulatory cell type when crossed with cell-type specific Cre-line (Fig. 4E,F), and does so with high specify as indicated by co-expression of GCaMP6f with each cell-type specific marker.

**Figure S5.**
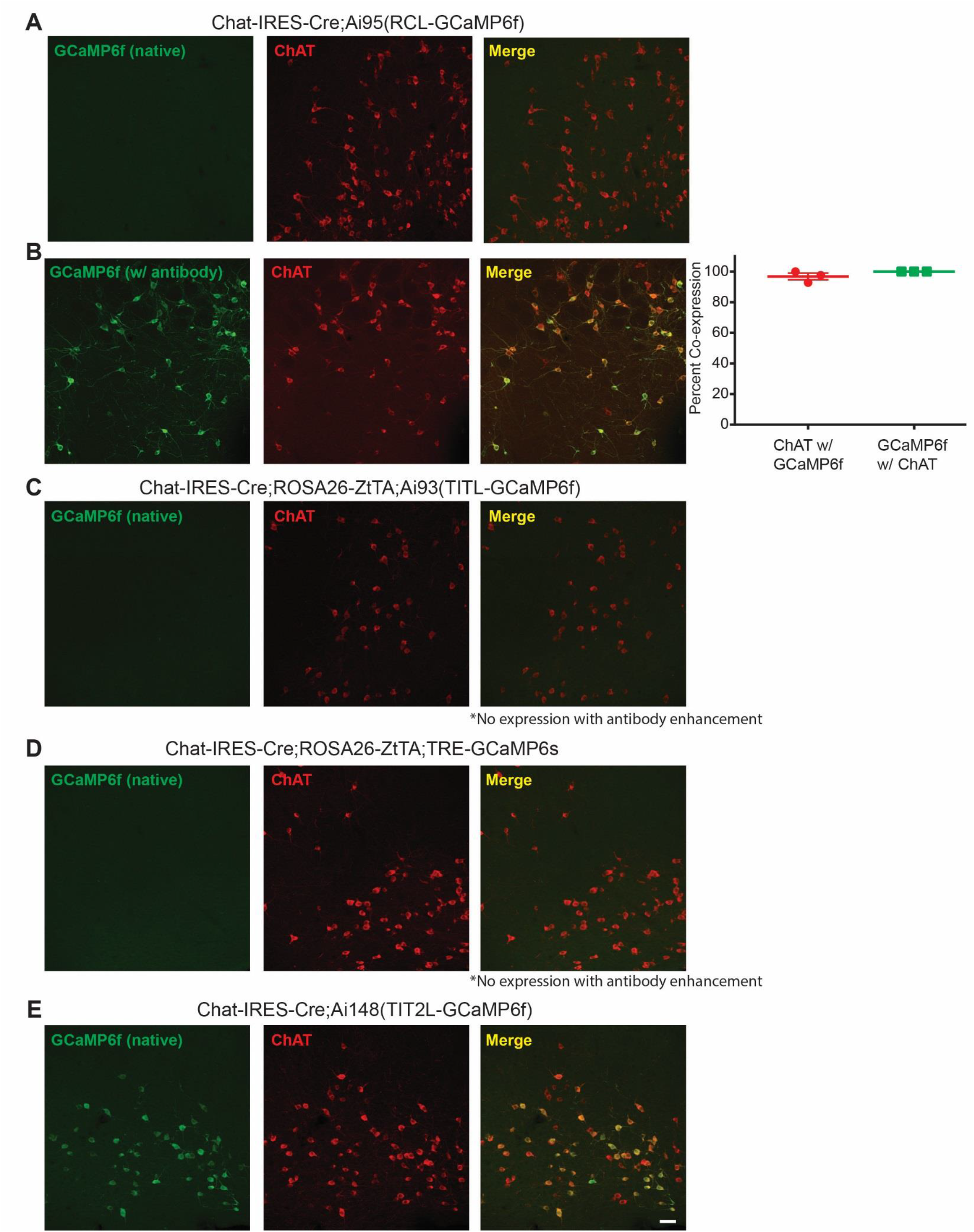
TIGRE2.0-based reporter line drives superior expression of GCaMP6f in basal forebrain cholinergic neurons compared to other reporters. Related to Figure 4. (**A-B**) Representative confocal images show that the Rosa26-based Cre reporter line Ai95 expresses GCaMP6f in cholinergic neurons when crossed to Chat-IRES-Cre mice, however, native GCaMP6f fluorescence in fixed sections is undetectable compared to TIGRE2.0 lines (compare panels A and E). In antibody-enhanced sections (**B**), Ai95-driven GCaMP6f expression is specific and widespread among cholinergic neurons as indicated by co-expression of GCaMP6f with choline acetyltransferase (ChAT)-positive neurons. Data are presented as mean ± S.E.M; n = 3 mice. (**C-D**) No GCaMP6f expression is observed in cholinergic neurons from crosses of Chat-IRES-Cre;Rosa26-ZtTA mice with either the TIGRE1.0 line, Ai93 (C) or a different TRE-GCaMP6s reporter (D) (Wekselblatt et al, 2016); n = 2-3 mice per genotype. * No expression can be detected even with immunofluorescence enhancement (data not shown). (**E**) The TIGRE2.0-based Ai148 reporter line drives expression of GCaMP6f in cholinergic neurons and at higher levels than previous GCaMP6 reporter lines. Scale bar: 50 μm.

**Figure S6.**
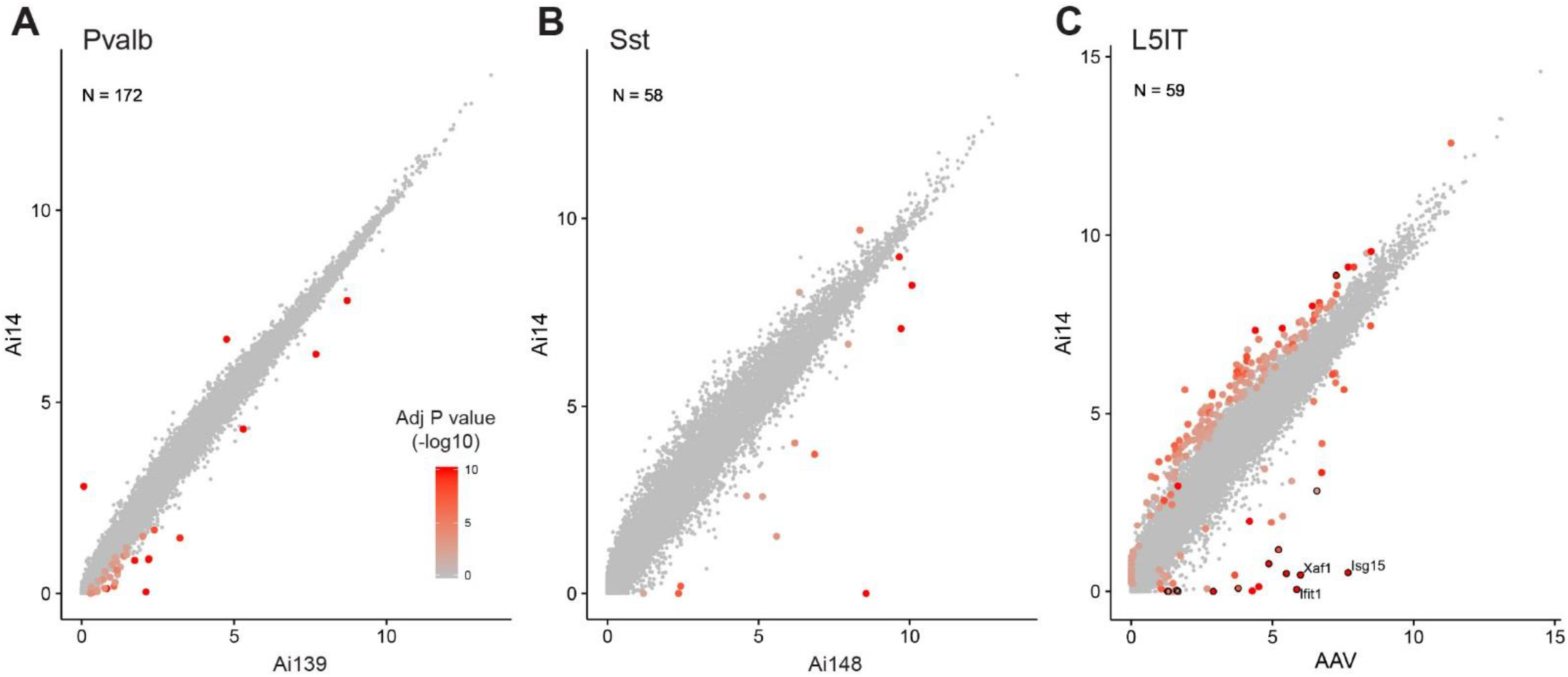
Global gene expression comparisons between TIGRE2.0-labeled or AAV-infected neurons and Ai14-labeled neurons by scRNA-seq. Related to Figure 4. (**A-B**) SMART-Seq v4 datasets from GFP+ inhibitory interneurons isolated from Pvalb-T2A-CreERT2;Ai139 (n = 172 cells, A) and Sst-IRES-Cre;Ai14;Ai148 (n = 58 cells, B) mice were mapped to reference transcriptomic cell types using nearest centroid classifiers. The same number of cells from Cre x Ai14 crosses with matching cell types and genders were chosen as control. Cells with similar cell types were then grouped into cell classes. We detected differentially expressed genes between TIGRE2.0 cells and corresponding Ai14 cells within the same cell class using limma voom mode. X and Y axes correspond to the average counts per million values in log scale, log2(cpm+1), for cells from Cre x TIGRE2.0 mice and Cre x Ai14 mice, respectively. Genes were colored based on −log10(adjusted P value) to indicate significant changes in expression levels. (**C**) SMART-Seq v4 datasets from GFP+ cells isolated from AAV-pCAG-FLEX-GFP-WPRE-pA virus injected Sim1-Cre_KJ18;Ai14 mice were mapped to reference transcriptomic cell types using nearest centroid classifiers. Cells were isolated 2 weeks after the AAV injection. The same number of non-AAV infected cells from the same Cre x Ai14 cross with matching cell types and genders were chosen as control. Cells belonging to related types were grouped into cell classes. There were n = 59 AAV-infected Sim1-Cre_KJ18;Ai14 cells mapped to the L5IT class. We detected differentially expressed genes between AAV-infected cells and corresponding cells from the same Cre x Ai14 cross without AAV injection within the same cell class using limma voom mode. X and Y axes correspond to the average counts per million values in log scale, log2(cpm+1), for AAV-infected cells and corresponding uninfected cells, respectively. Genes were colored based on - log10(adjusted P value) to indicate significant changes in expression levels. A number of genes upregulated in AAV-infected cells belong to a Type I interferon responsive group (GO: 0034340), and they are highlighted by black circles. The top 3 differentially expressed genes from this group were labeled by their names.

**Figure S7.**
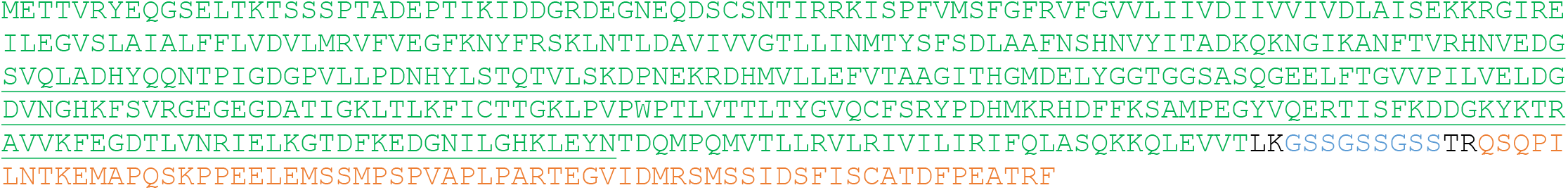
Protein sequence of soma-enriched voltage sensor ASAP2s-Kv. Related to Figure 5. The ASAP2s segment is labeled in green, with the circularly permuted GFP sequence underlined. The 65 amino acid segment from the C-terminus of the rat Kv2.1 potassium channel (orange) was fused to the C-terminus of ASAP2s via a flexible linker (blue). Black letters correspond to enzyme restriction sites. Distinct elements of the protein are color-coded. This segment is sufficient to target a heterologous protein to neuronal soma and proximal dendrites and has been repeatedly used for soma targeting of opsins.

**Figure S8.**
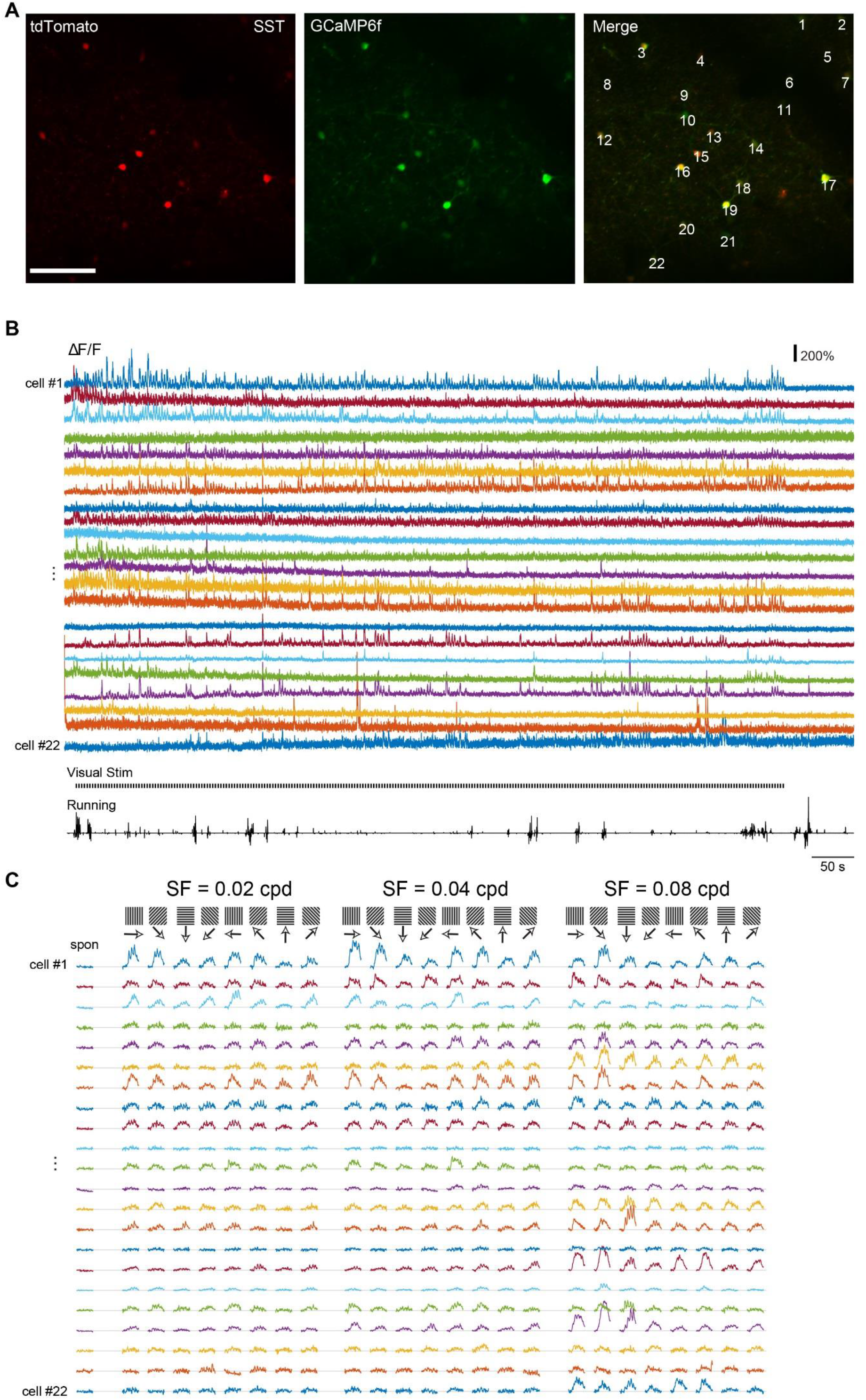
Strong and diverse visual responses of SST inhibitory interneurons reported by Ai148 mice. Related to Figure 6. (**A**) Two-photon calcium imaging Z-projection images (from time series) of the field-of-view as in Figure 6A showing red fluorescence (tdT, left), green fluorescence (GCaMP6f, middle) and merged (right) in a Sst-IRES-Cre;Ai14;Ai148 mouse. A total of 22 ROIs (representing individual cells, numbered) were analyzed for this field-of-view and results were shown in B and C. (**B**) ΔF/F traces for the 22 ROIs (cells) during the entire visual response mapping experiment. The mapping was performed with whole-screen sinusoidal drifting gratings showing 8 orientations (0° to 315° in 45° increments), 5 spatial frequencies (0.01, 0.02, 0.04, 0.08 and 0.16 cycle per degree) and 1 temporal frequency (2 Hz), presented at 80% contrast in a random sequence for 8 repetitions (STAR Methods). Running velocity is shown at the bottom. (**C**) Orientation tuning properties derived from the ΔF/F responses of individual SST cells.

**Figure S9.**
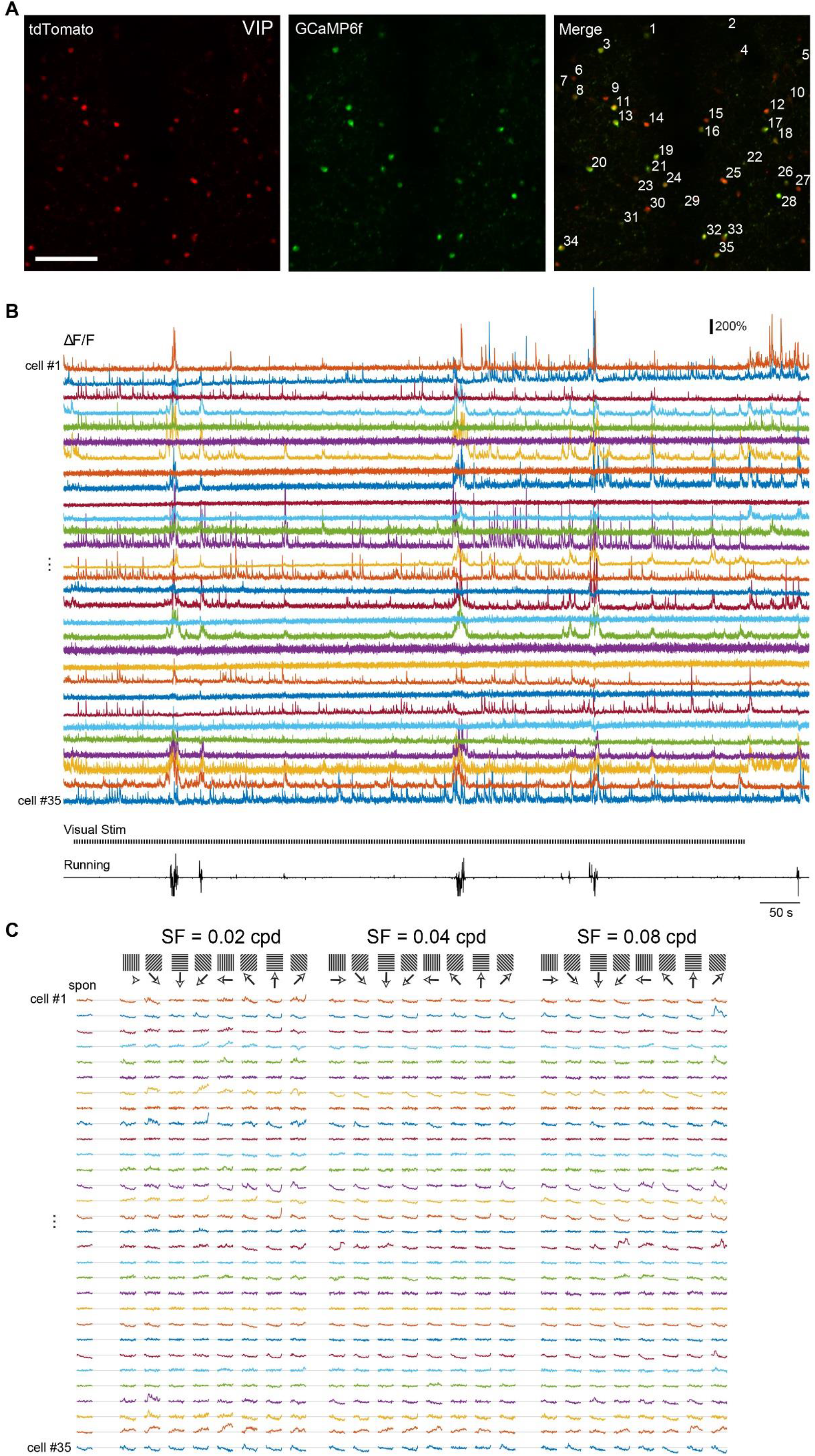
Strong and diverse visual responses of VIP inhibitory interneurons reported by Ai148 mice. Related to Figure 6. (**A**) Two-photon calcium imaging Z-projection images (from time series) of the field-of-view as in Figure 6B showing red fluorescence (tdT, left), green fluorescence (GCaMP6f, middle) and merged (right) in a Vip-IRES-Cre;Ai14;Ai148 mouse. A total of 35 ROIs (cells, numbered) were analyzed. (**B**) ΔF/F traces of the 35 ROIs (cells) during the entire visual response mapping. (**C**) Orientation tuning properties derived from the ΔF/F responses of individual VIP cells.

**Figure S10.**
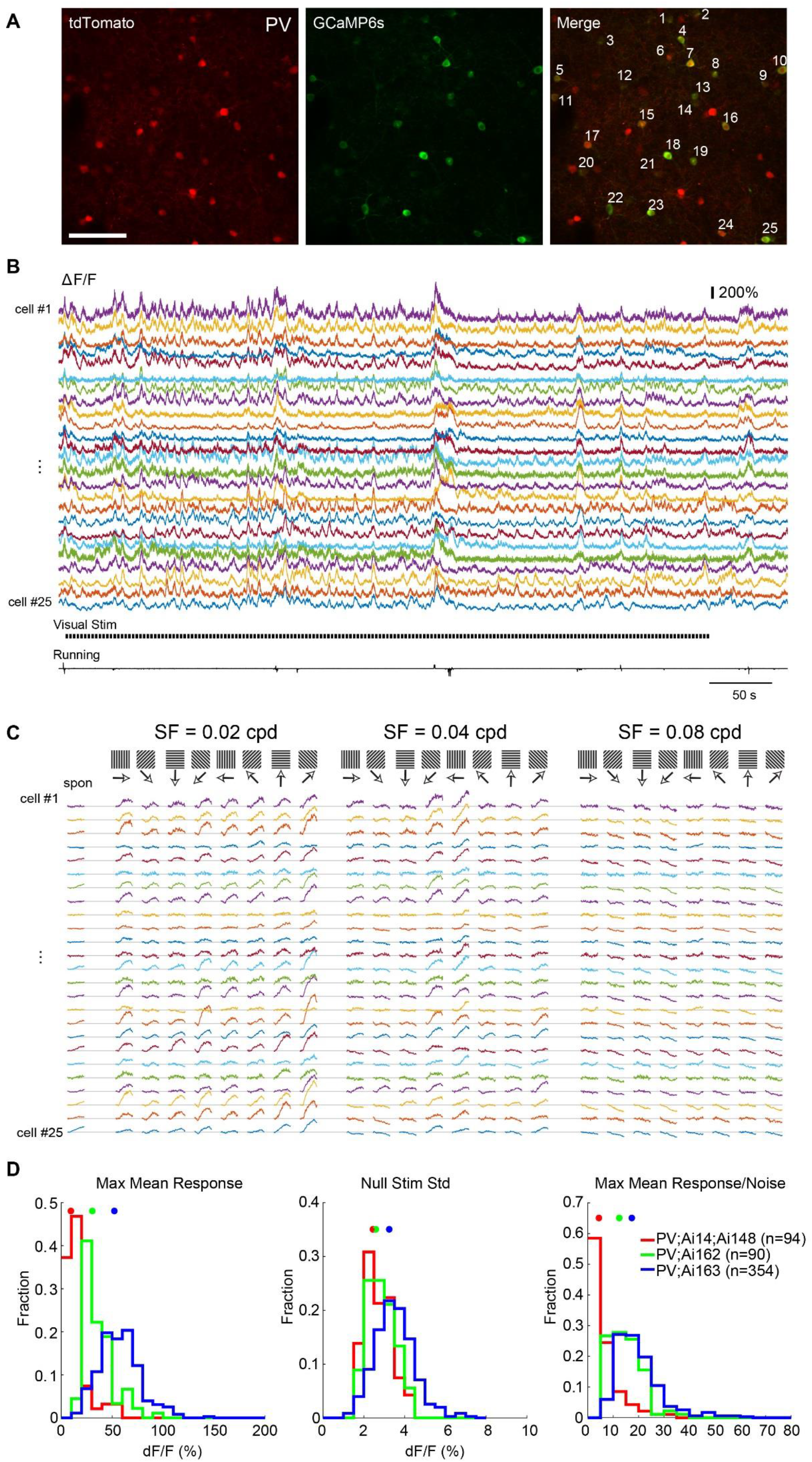
Strong and diverse visual responses of PV inhibitory interneurons reported by Ai163, Ai162 and Ai148 mice. Related to Figure 6. (**A**) Two-photon calcium imaging Z-projection images (from time series) of the field-of-view as in Figure 6C showing red fluorescence (tdT, left), green fluorescence (GCaMP6f, middle) and merged (right) in a Pvalb-IRES-Cre;Ai163 mouse. A total of 25 ROIs (cells, numbered) were analyzed. (**B**) ΔF/F traces of the 25 ROIs (cells) during the entire visual response mapping. (**C**) Orientation tuning properties derived from the ΔF/F responses of individual PV cells. (**D**) Improved expression and functionality of GECI in PV cells from Ai148 to Ai162 and Ai163, as shown in a comparison among Pvalb-IRES-Cre;Ai14;Ai148 (red), Pvalb-IRES-Cre;Ai162 (green) and Pvalb-IRES-Cre;Ai163 (blue) mice. Left: Population histogram of the mean max responses. Middle: Population histogram of the standard deviation of ΔF/F during gray screens (null stimuli). Right: Population histogram of the estimated signal-to-noise ratio. Scale bar: 100 μm.

**Figure S11.**
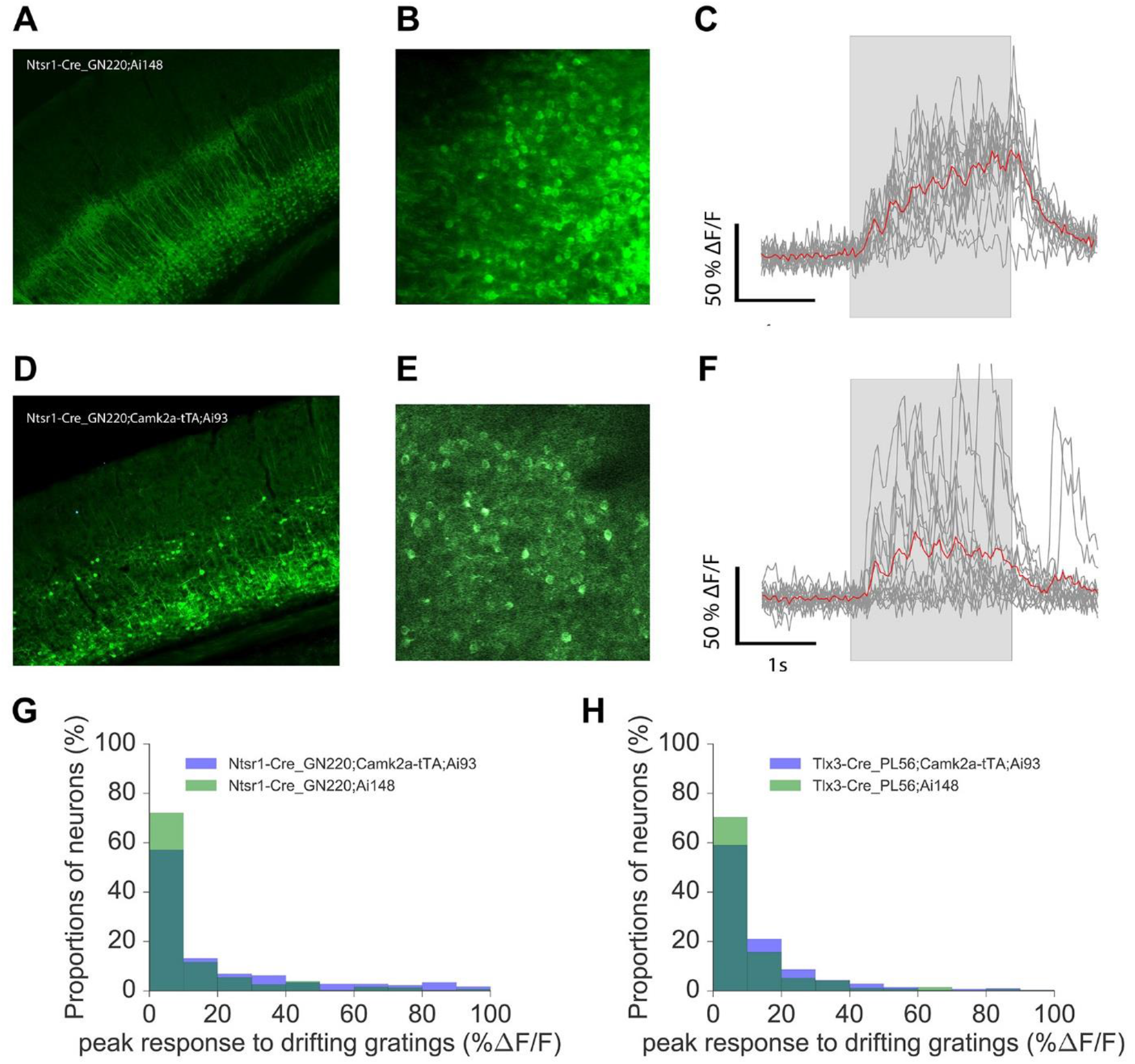
Comparison of visually evoked calcium responses in deep cortical layers in Ai148 and Ai93 mice. Related to Figure 6. (**A**) A representative coronal section in VISp of an Ntsr1-Cre_GN220;Ai148 mouse showing GCaMP6f expression in layer 6 neurons specifically. (**B**) A representative averaged two-photon image (of 500 frames) recorded *in vivo* in VISp layer 6 of the mouse shown in A. (**C**) Example neuronal responses to drifting gratings for a cell recorded in B. The red trace represents the average response. (**D**) A representative coronal section in VISp of an Ntsr1-Cre_GN220;Camk2a-tTA;Ai93 mouse showing GCaMP6f expression in layer 6 neurons as well as some upper layer neurons. (**E**) A representative averaged two-photon image (of 500 frames) recorded *in vivo* in VISp layer 6 of the mouse shown in D. (**F**) Example neuronal responses to drifting gratings for a cell recorded in E. The red trace represents the average response. (**G**) Distribution of peak amplitudes of visually evoked responses to drifting gratings for all cells recorded in Ntsr1-Cre_GN220 mice (total of 183 cells in Ntsr1-Cre_GN220;Camk2a-tTA;Ai93, and 310 cells in Ntsr1-Cre_GN220;Ai148 mice). (**H**) Distribution of peak amplitudes of visually evoked responses to drifting gratings for all cells recorded in another Cre line, Tlx3-Cre_PL56, which drives GCaMP6f expression in cortical layer 5 neurons (total of 281 cells in Tlx3-Cre_PL56;Camk2a-tTA;Ai93, and 349 cells in Tlx3-Cre_PL56;Ai148 mice).

**Figure S12.**
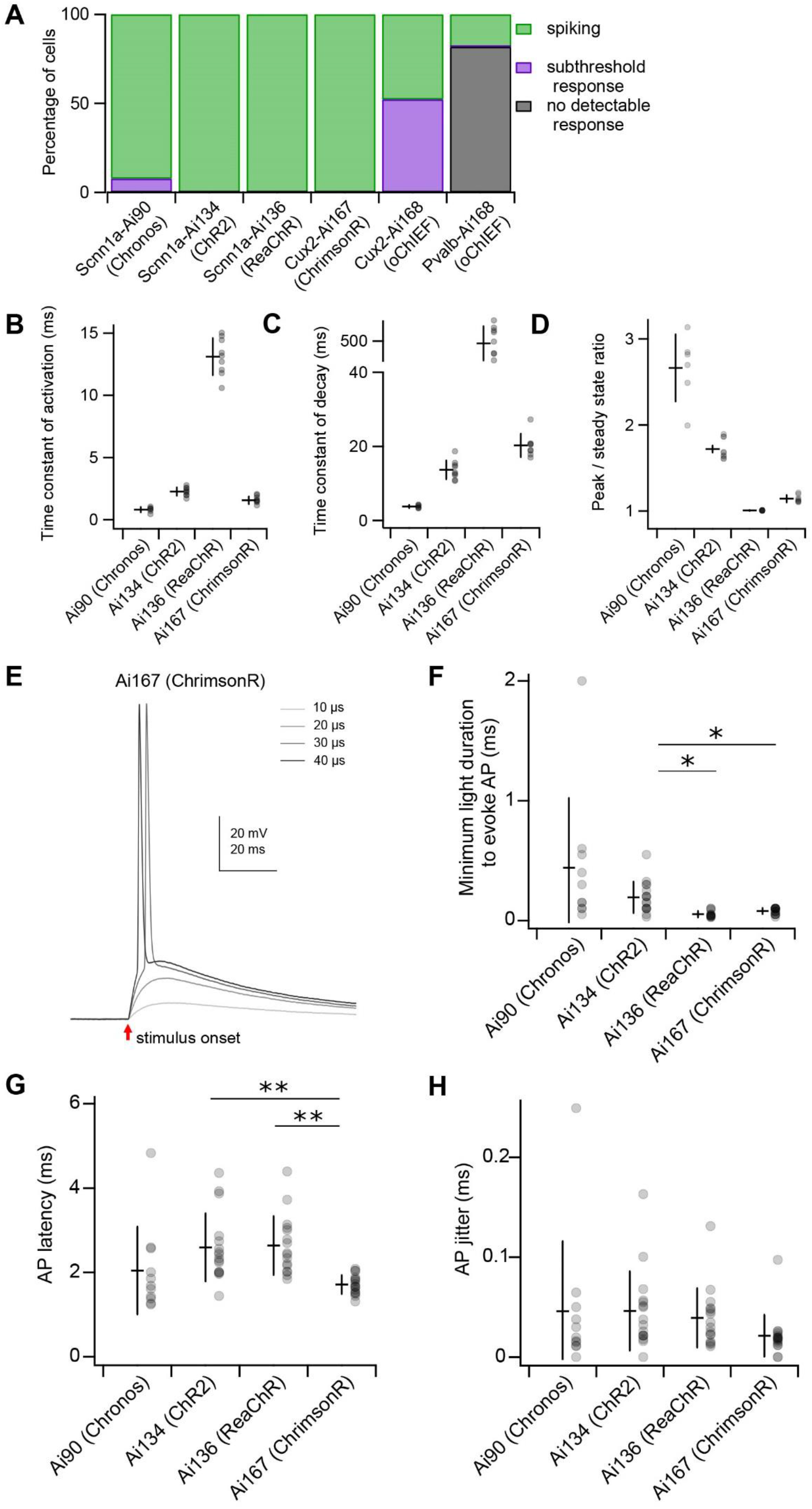
Characterization of light-evoked responses for Ai90, Ai134, Ai136, Ai167 and Ai168 transgenic lines. Related to Figure 7. In addition to mouse strains described in Figure 7, also examined are Cux2-CreERT2;Ai168 and Pvalb-IRES-Cre;Ai168 mice expressing oChIEF in cortical layer 2/3/4 excitatory neurons and PV+ interneurons, respectively. For all data presented here, neurons recorded from Ai90, Ai134 and Ai168 animals were stimulated with 470 nm light, whereas experiments characterizing Ai136 and Ai167 mice used 590 nm light. (**A**) Percentage of cells in each transgenic line displaying action potential (AP) firing, subthreshold response or no response to optical stimulation. We first tested the ability of each neuron to generate light-evoked APs with 1 ms light pulses. If AP firing was not observed, light exposure was prolonged (up to 50 ms) to further test the sensitivity of the cell to optical stimulation. Except for the Ai168 lines, nearly all cells fired in response to a 1 ms light pulse (see panel **F**). (**B**) Time constants of activation measured by a mono-exponential fit from the onset of the light pulse to the peak of the photocurrent (see Fig. 7A). Ai90: 0.81 ± 0.21 ms; Ai134: 2.3 ± 0.3 ms; Ai136: 13.1 ± 1.5 ms; Ai167: 1.6 ± 0.3 ms. (**C**) Time constants of deactivation measured by a mono-exponential fit from the termination of the light pulse to the return of the current to baseline (see Fig. 7A). Ai90: 3.8 ± 0.4 ms; Ai134: 13.7 ± 2.5 ms; Ai136: 478 ± 166 ms; Ai167: 20.2 ± 3.1 ms. (**D**) Peak/steady-state ratio. Steady-state was measured as the average current amplitude during the final 5 ms of the 100 ms light exposure (see Fig. 7A). Ai90: 2.66 ± 0.39; Ai134: 1.72 ± 0.04; Ai136: 1.01 ± 0.003; Ai167: 1.14 ± 0.04 (*n* = 6-9 cells for each group in panels B-D). Measures of photocurrent kinetics were significantly different between all groups. For time constant of activation: *P* < 0.0005; time constant of decay: *P* < 0.0005; peak/steady state ratio: *P* < 0.005. (**E**) Representative responses of a Cux2-CreERT2;Ai167 cell to saturating light pulses of increasing duration. (**F**) Minimum light durations required to evoke an AP for each transgenic line. Minimum duration in Ai90: 0.44 ± 0.58 ms; Ai134: 0.19 ± 0.13 ms; Ai136: 0.052 ± 0.027 ms; Ai167: 0.078 ± 0.025 ms. (**G**) Latency of spiking in response to a 1 ms saturating light pulse. Ai90: 2.0 ± 1.0 ms; Ai134: 2.6 ± 0.8 ms; Ai136: 2.6 ± 0.7 ms; Ai167: 1.7 ± 0.2 ms. (**H**) Jitter of spiking in response to a 1 ms saturating light pulse. Ai90: 46 ± 70 μs; Ai134: 46 ± 40 μs; Ai136: 39 ± 30 μs; Ai167 21 ± 21 μs. All data within this figure are plotted as the mean ± standard deviation. \**P* < 0.001; \*\**P* < 0.0001. *n* = 10-16 cells per transgenic line.

**Figure S13.**
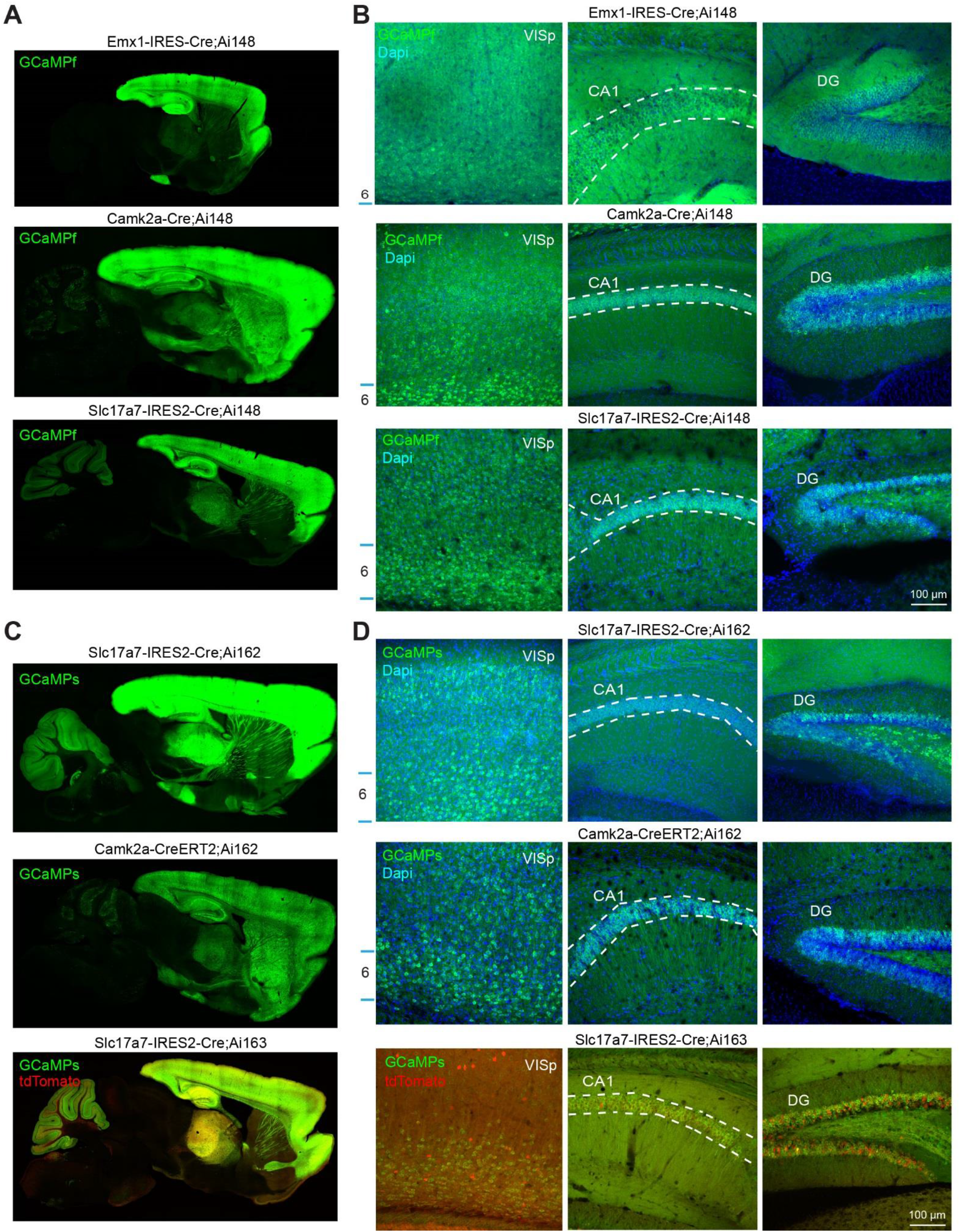
Adverse effects associated with strong and broad transgene expression. (**A-D**) Representative confocal images of native GCaMP6f, GCaMP6s, or tdTomato fluorescence from Ai148 (A-B), Ai162 and Ai163 (C-D) mice crossed to various cortical pan-excitatory Cre lines. Expression throughout the brain, in the primary visual cortex (VISp) and within hippocampal subfields (CA1 and dentate gyrus) is shown. The approximate location of layer 6 within the cortex and the pyramidal layer of CA1 (between dashed lines) are annotated in the 20X images. All montage and 20X images were acquired using the same instrument settings.

**Table S1.**
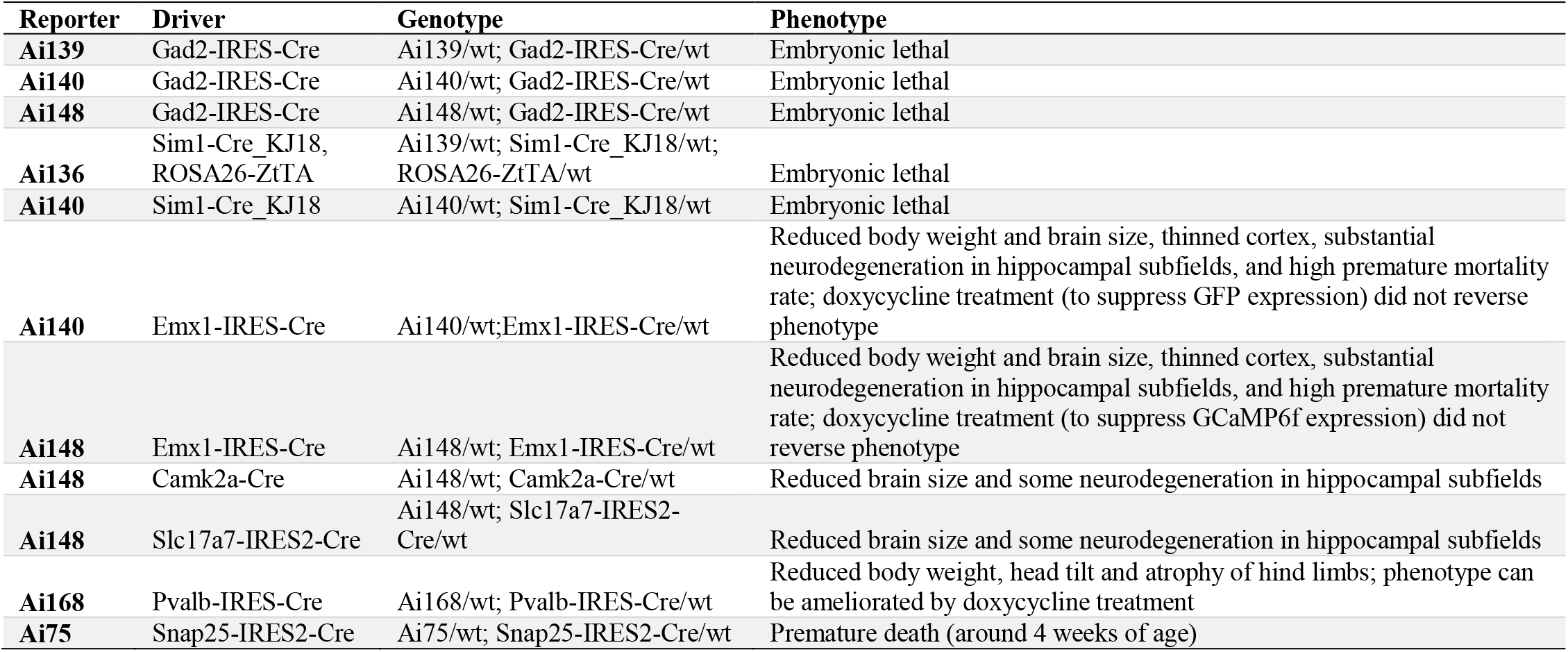
Problematic reporter and driver line crosses.

**Figure S14.**
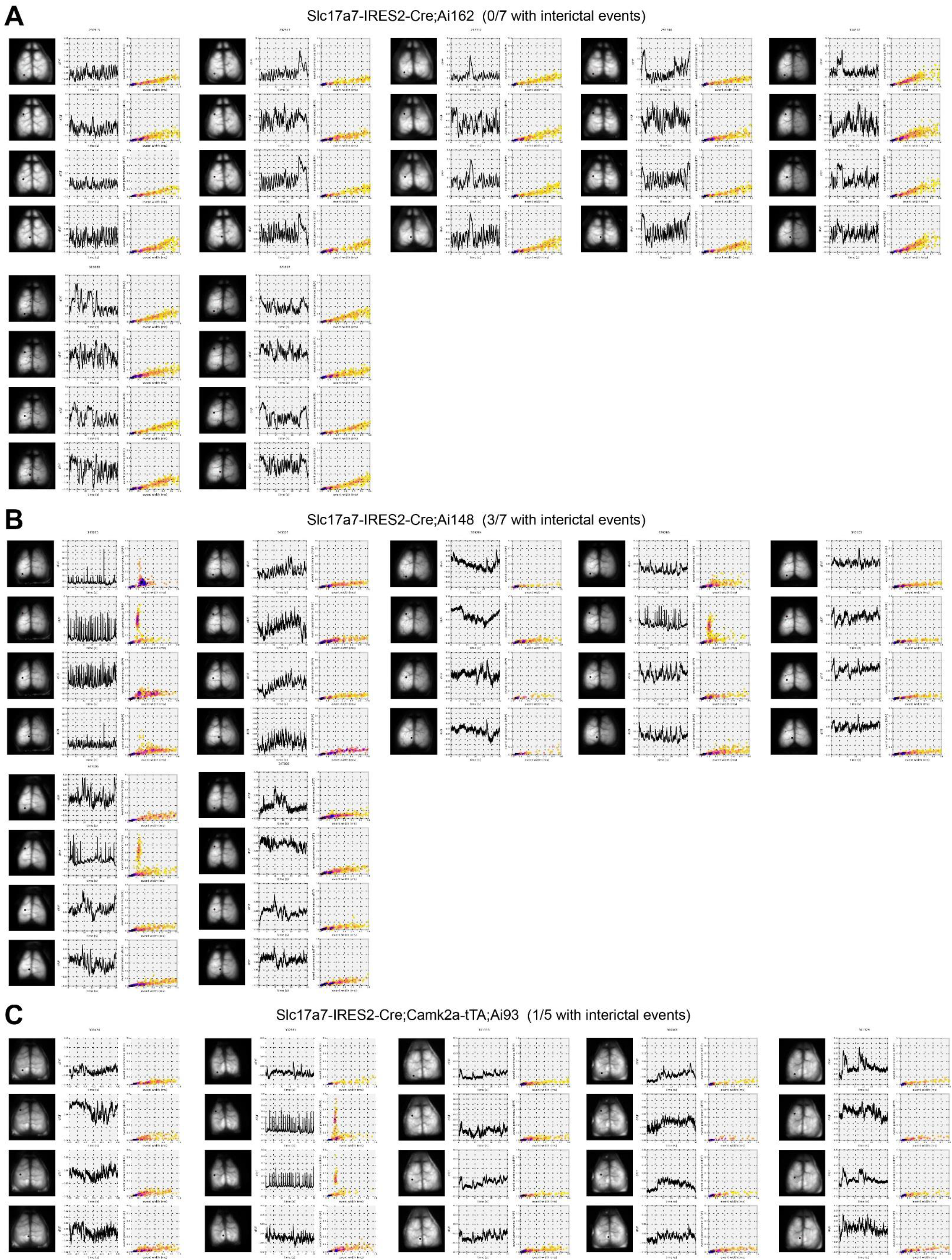
All observations of normal and aberrant cortical activity in cortical pan-excitatory GCaMP6 expressing lines. Related to Figure 8. (**A**) Wide-field calcium imaging of 7 Slc17a7-IRES2-Cre;Ai162 mice. For each mouse, GCaMP6 fluorescence was quantified from four cortical areas (from top down): primary visual cortex, primary motor cortex, primary somatosensory cortex, and retrosplenial cortex (same for B and C). None of the 7 mice examined had interictal activity. (**B**) Wide-field calcium imaging of 7 Slc17a7-IRES2-Cre;Ai148 mice. We detected interictal activity in 3 of the 7 mice examined. (**C**) Wide-field calcium imaging of 5 Slc17a7-IRES2-Cre;Camk2a-tTA;Ai93 mice. We detected interictal activity in 1 of the 5 mice examined.

**Table S2.** Animals from Fig. S14. Animals with aberrant activity are highlighted (gray), observations with aberrant activity are highlighted in red, with the corresponding sex and age of animals at the time of each observation. The observations plotted in Fig. S14 are indicated in bold.

| Genotype | Mouse ID# | Sex | Age at observation<br>(postnatal days) |
| --- | --- | --- | --- |
| Slc17a7-IRES2-Cre;Ai162 | 292915 | F | **52** |
| Slc17a7-IRES2-Cre;Ai162 | 292917 | F | **52** |
| Slc17a7-IRES2-Cre;Ai162 | 297312 | M | **110** |
| Slc17a7-IRES2-Cre;Ai162 | 297310 | M | **110** |
| Slc17a7-IRES2-Cre;Ai162 | 304572 | F | **170** |
| Slc17a7-IRES2-Cre;Ai162 | 315689 | F | **120** |
| Slc17a7-IRES2-Cre;Ai162 | 321607 | F | **97, 191** |
| Slc17a7-IRES2-Cre;Ai148 | 343025 | M | **77** |
| Slc17a7-IRES2-Cre;Ai148 | 343027 | F | **77, 99** |
| Slc17a7-IRES2-Cre;Ai148 | 329284 | M | **67** |
| Slc17a7-IRES2-Cre;Ai148 | 329286 | M | **67** |
| Slc17a7-IRES2-Cre;Ai148 | 347103 | M | **57, 79** |
| Slc17a7-IRES2-Cre;Ai148 | 347105 | F | **71** |
| Slc17a7-IRES2-Cre;Ai148 | 347893 | M | **67, 75** |
| Slc17a7-IRES2-Cre;Camk2a-tTA;Ai93 | 303474 | F | **69, 118** |
| Slc17a7-IRES2-Cre;Camk2a-tTA;Ai93 | 302981 | M | 60, **68, 111** |
| Slc17a7-IRES2-Cre;Camk2a-tTA;Ai93 | 301333 | F | **67, 118** |
| Slc17a7-IRES2-Cre;Camk2a-tTA;Ai93 | 306069 | F | **67, 96** |
| Slc17a7-IRES2-Cre;Camk2a-tTA;Ai93 | 301329 | M | **67, 118** |

